# Dynamical control enables the formation of demixed biomolecular condensates

**DOI:** 10.1101/2023.01.04.522702

**Authors:** Andrew Z. Lin, Kiersten M. Ruff, Ameya Jalihal, Furqan Dar, Matthew R. King, Jared M. Lalmansingh, Ammon E. Posey, Ian Seim, Amy S. Gladfelter, Rohit V. Pappu

## Abstract

Macromolecular phase separation underlies the regulated formation and dissolution of biomolecular condensates. What is unclear is how condensates of distinct and shared macromolecular compositions form and coexist within cellular milieus. Here, we use theory and computation to establish thermodynamic criteria that must be satisfied to achieve compositionally distinct condensates. We applied these criteria to an archetypal ribonucleoprotein condensate and discovered that demixing into distinct protein-RNA condensates cannot be the result of purely thermodynamic considerations. Instead, demixed, compositionally distinct condensates arise due to asynchronies in timescales that emerge from differences in long-lived protein-RNA and RNA-RNA crosslinks. This type of dynamical control is also found to be active in live cells whereby asynchronous production of molecules is required for realizing demixed protein-RNA condensates. We find that interactions that exert dynamical control provide a versatile and generalizable way to influence the compositions of coexisting condensates in live cells.

## Main Text

Biomolecular condensates arise via spontaneous and driven phase transitions of complex mixtures of multivalent protein and RNA molecules ^1–5^. A key challenge is understanding how condensates with shared and distinct macromolecular compositions can coexist as distinct functional entities ^6^ (**Figure 1a**). The importance of this issue is emphasized by observations in the filamentous fungus *Ashbya gossypii* where Langdon et al., ^7^ reported the presence of coexisting ribonucleoprotein condensates that share the protein Whi3. Specifically, the cyclin RNA *CLN3* forms a distinct condensate with Whi3 that does not colocalize with condensates formed by Whi3 with *BNI1* or *SPA2* ^7^, which are RNA molecules encoding proteins that regulate actin. These compositionally distinct condensates function in separate cell processes^8–10^, controlling nuclear division and cell polarity. Previous studies showed that protein-protein, RNA-protein, and RNA-RNA interactions contribute to the formation and identity of Whi3 condensates. However, the interactions that drive phase separation, specifically the driving forces for demixing that generate compositionally distinct condensates remain unclear ^8,9^.

**Figure 1:**
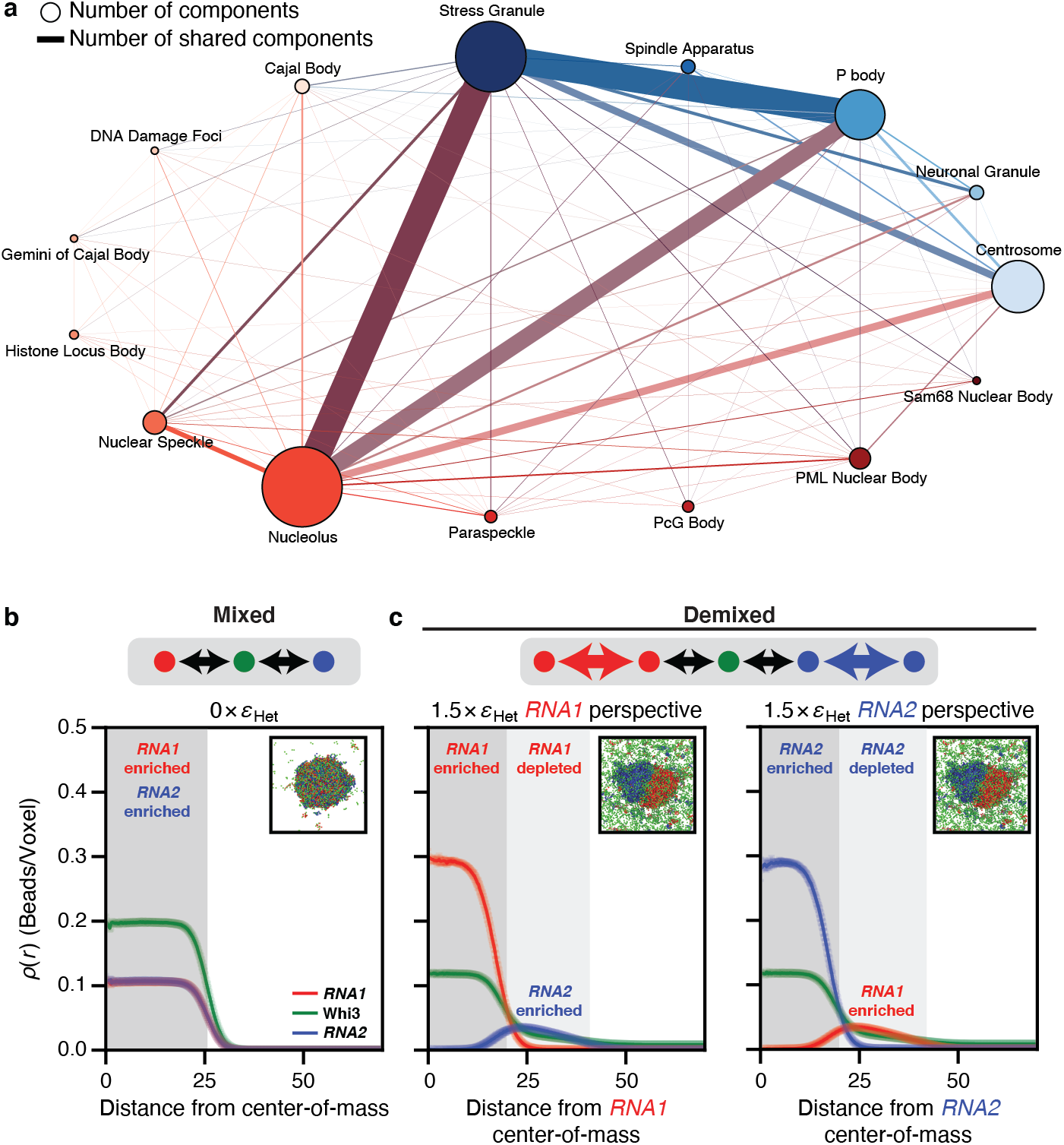
Simulations show that the interplay between homotypic and heterotypic interactions modulates the formation of mixed versus demixed phases. (a) Graph of components in condensates in cells from *Homo sapiens* as extracted from DrLLPS ^14^. Here, the node size correlates with the number of components within the condensate and the edge width correlates with the number of shared components between two condensates. Each nuclear condensate is depicted as a node of a distinct size and reddish hue. Likewise, each cytoplasmic condensate is depicted as a node of a distinct size and blueish hue. Edges between nodes are shown in the color of the mixture of the two node colors. (b-c) Radial density profiles from LaSSI lattice-based simulations of a ternary system designated as *RNA1* + *RNA2* + a Whi3 mimic (see Methods). Black arrows denote heterotypic interactions, whereas colored arrows denote homotypic interactions. (b) The system forms a well-mixed condensate when the RNAs engage in purely heterotypic interactions with the Whi3 mimic. (c) In contrast, demixed condensates are formed when the RNAs have strong homotypic interactions as well. Here, εHet = −2*k_B_T* refers to the strength of RNA interactions with the Whi3 mimic. The right two plots show results from the same simulations. In the middle plot, the macromolecular densities are quantified from the perspective of the center-of-mass of *RNA1* whereas in the plot on the right, the density profiles are plotted from the perspective of the center-of-mass of *RNA2*.

The formation of a single condensate versus sets of distinct coexisting condensates can be predicted by mean field theories where the relevant parameters are the variance of intermolecular interaction strengths and the numbers of components in multicomponent systems ^11^. In ternary systems, such as mixtures of Whi3 with *CLN3* and *BNI1* in an aqueous buffer, the relevant considerations are the interplay of solvent-mediated homotypic and heterotypic interactions (**Figure S1**). We used computations to ascertain the types of interactions that must be encoded to ensure that a protein can spontaneously form distinct condensates with two different types of RNA molecules. We performed lattice-based coarse-grained simulations ^12,13^ of a model system in which two high valency linear polymers (designated as *RNA1* and *RNA2)* interact with the same low valency linear polymer (a Whi3 mimic) with varying types of interactions among *RNA1, RNA2*, and the Whi3 mimic. Here, valency refers to the numbers of cohesive motifs (stickers) that can engage in complementary homotypic or heterotypic interactions. When the effective interactions of *RNA1* and *RNA2* with the Whi3 mimic are the same in strength and purely heterotypic in nature, then two coexisting phases form, with the dense phase encompassing all three macromolecules (**Figure 1b**). In contrast, we observe a demixing of dense phases when *RNA1* and *RNA2* have favorable homotypic interactions of strengths that are equivalent to or stronger than heterotypic interactions with the Whi3 mimic (**Figure 1c**). These dense phases feature *RNA1*-rich and *RNA2-* rich territories, with the Whi3 mimic being dispersed equivalently in both regions (**Figure 1c, Figures S2-3**). Other interaction modes can also engender the demixing phenotype. These include repulsions between different RNAs (**Figures S4-5**), which we deem unlikely to be relevant since electrostatic repulsions cannot be selectively imposed among different RNA molecules. Strongly asymmetric interactions featuring heterotypic interactions of very different strengths between the RNAs and Whi3 mimics can also enable demixing (**Figures S6-7**).

Taken together, the simulations show that assessments of relative contributions of homotypic versus heterotypic interactions are critical for extracting the relevant thermodynamic driving forces of the phase behaviors of Whi3-RNA mixtures. Accordingly, we set out to use *in vitro* measurements to assess the contributions of homotypic versus heterotypic interactions and determine if there are significant asymmetries in heterotypic interactions.

We first developed a method to extract the relative strengths and contributions of homotypic and heterotypic interaction to phase behaviors of mixtures. For this, we analyzed the shapes of phase boundaries in phase diagrams by bootstrapping against results from coarse-grained simulations of binary mixtures of RNA and Whi3 mimics ^4,12^. **Figure 2a** shows the phase boundaries plotted in terms of the concentrations of molecules when the system features heterotypic interactions between RNA and Whi3 mimics with or without equivalent homotypic interactions of the RNA mimic. For purely heterotypic interactions the phase boundary forms a closed loop ^2,12,15,16^. In contrast, when homotypic interactions of the RNA mimic are equivalent to heterotypic interactions with the Whi3 mimic, the RNA mimic can phase separate on its own leading to a flattening of the dilute arm of the phase boundary with decreased concentration of the Whi3 mimic (**Figure 2a**).

**Figure 2:**
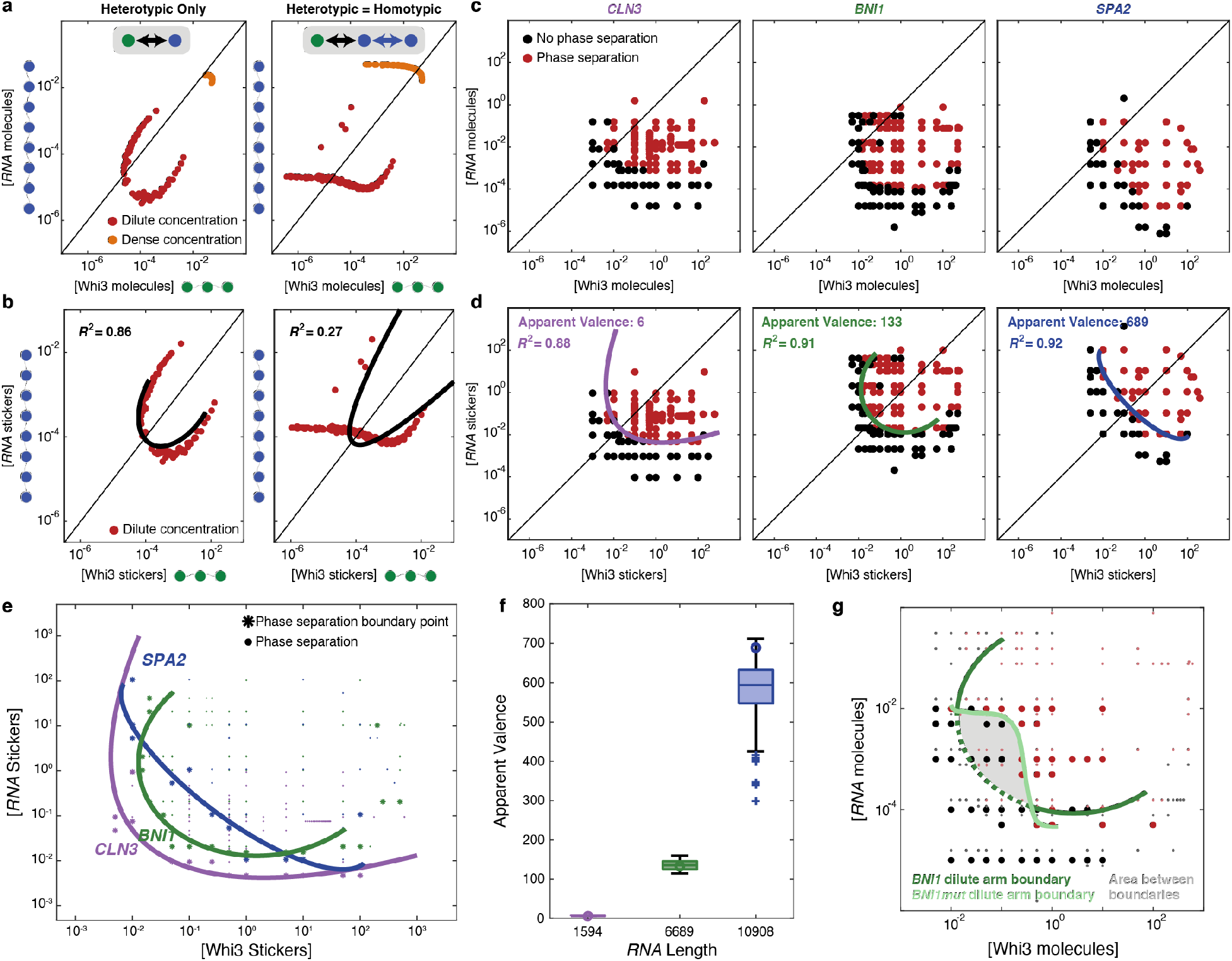
Phase boundaries for two-component systems can be used to extract the relative interaction strengths of homotypic and heterotypic interactions and the apparent valence of the RNAs. (a) Phase boundaries computed using LaSSI lattice-based simulations for a system with RNA and Whi3 mimics when interactions are purely heterotypic (left) or homotypic interactions are included among RNA molecules (right). Here, the RNA and Whi3 mimics have sticker valencies of eight and three, respectively. (b) The phase boundaries in panel (a) are recast in terms of concentrations of stickers. Black lines denote elliptical fits and the *R^2^* values quantify the goodness of the fit. (c) Phase boundaries, measured *in vitro*, of Whi3 with *CLN3* (left), *BNI1* (center), and *SPA2* (right) determined by confocal microscopy. Here, the phase boundaries are depicted in terms of concentrations of macromolecules. (d) The data in panel (c) are recast in terms of concentrations of stickers (see Methods). Elliptical fits are shown in solid color lines and the *R*^2^ values quantify the goodness of the fit. (e) The derived phase boundaries, recast in terms of concentrations of protein and RNA stickers are shown for each of the three Whi3-RNA systems. Here, colored points represent concentrations at which phase separation was observed by confocal microscopy. Larger points denote the points that were used to generate consensus elliptical boundaries for each of the three systems. (f) Plot of the inferred apparent valence of stickers for each of the RNA molecules (see Methods). Circles denote the apparent valence that corresponds to the minimum overlap between the one- and two-phase regimes. (g) Measured phase boundaries for Whi3-*BNI1* and *Whi3-BNI1mut* mixtures. Here, black, and dark red points denote points in the one-versus two-phase regime for the *Whi3-BNI1mut* system, whereas grey and light red points denote points within the one-versus two-phase regime for the Whi3-*BNI1* system. A dark green ellipse is fit to the measured phase boundary for the Whi3-*BNI1* system, whereas the light curve denotes the low concentration arm of the measured phase boundary of the Whi3-*BNI1mut* system. Scrambling the cognate Whi3-binding sites on *BNI1* eliminates phase separation in low concentration regime (sub-micromolar, grey area) for Whi3 and *BNI1mut* while preserving phase separation through the rest of the concentration regimes. All concentrations for *in vitro* measurements are in units of μM.

For phase separation driven by dominant heterotypic interactions the phase boundary should be symmetrical about the diagonal ^2,12,15–18^. While this is trivially true of molecules with equivalent numbers of cohesive motifs, known as stickers, it is also true if the titrations along each of the axes quantify the concentrations of stickers rather than molecules ^16^. For the case of purely heterotypic interactions, phase separation should be most favored when sticker concentrations are balanced between the two molecules. **Figure 2b** shows a remapping of phase boundaries in terms of sticker concentrations. For this, we leveraged the fact that we have *a priori* knowledge of the numbers of stickers in each of the multivalent polymers. We then quantified the degree of symmetry about the diagonal by fitting an ellipse to the dilute arm of the measured and mirrored phase boundary (see methods). If the phase boundary is symmetrical about the diagonal, as expected in a system driven by heterotypic interactions, then a single ellipse is an optimal locus of points along the phase boundary. The addition of homotypic interactions degrades the ability to symmetrize about the diagonal leading to a poorer fit of a single ellipse (**Figure 2b**). Therefore, remapping phase boundaries in terms of concentrations of stickers, and quantifying the degree to which the dilute concentration arm can be described by a single ellipse, can help in discerning the interplay between heterotypic and homotypic interactions.

Next, we applied insights from our computations to interpret measured phase boundaries of distinct binary mixtures *in vitro*. **Figure 2c** shows the measured phase boundaries for Whi3 with each of three different cognate RNA molecules, *viz., CLN3, BNI1*, and *SPA2*. Qualitatively, the phase boundaries do not show signatures of dominant homotypic interactions for Whi3 or any of the RNAs. To rigorously test this hypothesis, we determined the degree to which measured phase boundaries could be symmetrized about the diagonal, assessed fit of a single ellipse, and estimated apparent valence in the system.

To assess if phase boundaries can be symmetrized about the diagonal, the phase boundaries were remapped in terms of sticker concentrations. Stickers can enable homotypic and heterotypic interactions ^19^. The glutamine-rich region and the RRM of Whi3 are essential for driving phase separation. Of these, the RRM is known to engage in site-specific interactions with the cognate RNA molecules. For our analysis of measured phase boundaries, the null hypothesis is that the phase behavior is driven primarily by heterotypic interactions. Accordingly, we set the valence of RNA-binding stickers on Whi3 to be one. This assumption simplifies the analysis where the quantity to be extracted is the ratio of the numbers of stickers on the RNA molecules to the numbers of stickers on the Whi3 protein. Our choice makes the apparent valence (apparent number) of Whi3-binding stickers on each of the RNA molecules the titratable free parameter. We vary this parameter and quantify the extent to which measured phase boundaries can be symmetrized about the diagonal (Methods, **Figure S8**).

The measured phase boundaries, remapped onto sticker concentrations, can be well-described by system-specific closed loops, with *R^2^* values of 0.88-0.92 supporting strong fit to a single ellipse (**Figure 2d**). This suggests that heterotypic interactions are the main drivers of Whi3-RNA phase separation. As shown in **Figure 2b**, the symmetrization would have failed if homotypic interactions were on an equal footing with heterotypic interactions. Our findings do not imply the absence of homotypic interactions. Instead, they imply that, on balance, the contributions of heterotypic interactions are dominant over any homotypic protein-protein and RNA-RNA interactions, especially along directions that are parallel to the diagonal in a plane defined by concentrations of Whi3 and RNA stickers.

The numbers of cognate-binding sites for Whi3 are 5 each for *CLN3* and *BNI1* and 11 for *SPA2*. However, analysis of the measured phase boundaries leads to the inference that *CLN3, BNI1*, and *SPA2* have apparent sticker valencies of 6, 133, 689, respectively (**Figure 2f**). This suggests that there are additional non-cognate stickers within each of the RNA molecules. The numbers of non-cognate sites increase with length, and the ratio of non-cognate to cognate stickers is 0.2, 25.6, and 61.6 for *CLN3, BNI1*, and *SPA2*, respectively. It is noteworthy that *BNI1* and *SPA2*, which both share an enrichment of non-cognate sites, colocalize to the same Whi3 condensates in cells and contribute to the same function ^7^.

If there are non-cognate sites within longer RNA molecules, then abrogation of cognate sites should still preserve phase separation. To test for this, we measured *in vitro* phase boundaries for the two-component system comprising Whi3 and a *BNI1* mutant (*BNI1mut*) in which the cognate Whi3-binding sites were disrupted by scrambling the consensus Whi3-binding motif. The measured phase boundary for the *Whi3-BNI1mut* system overlaps with that of the Whi3-*BNI1* system at high Whi3 and low RNA concentrations and vice versa (**Figure 2g**). Thus, while noncognate sites can drive phase separation, their interactions are weaker than those of cognate Whi3-binding sites of *BNI1*. This inference comes from measured phase boundaries where the combination of Whi3 and *BNI1* concentrations that lie between the dashed and solid curve in **Figure 2g** defines the regime where phase separation requires cognate Whi3-binding sites of *BNI1*. The concave cutout on the measured phase boundary of Whi3 with *BNI1mut* comprises sections that are roughly horizontal and vertical (see light green boundary in **Figure 2g**). These sections represent protein and RNA concentrations where homotypic interactions become important contributors to phase separation. In mixtures of Whi3 with wild-type *BNI1* this region of the phase diagram is masked by the dominance of heterotypic interactions due to the cognate binding sites.

Our results suggest that heterotypic interactions between Whi3 and RNA molecules are the main drivers of phase separation that give rise to Whi3-RNA condensates in binary mixtures. Heterotypic interactions involve Whi3 and its cognate RNA-binding sites, as well as additional non-cognate sites of unknown sequence specificity. Therefore, the demixing phenotypes observed in cells are unlikely to be due to the dominance of homotypic interactions over heterotypic ones. Further, for demixing to be thermodynamically driven in systems where the phase behavior is dominated by heterotypic interactions, the driving forces for phase separation between the different binary mixtures must be significantly different (**Figure S1, S6-7**) ^20^. We find that the measured phase boundaries generally show less than a 4-fold difference between Whi3-*CLN3* and Whi3-*BNI1* binary mixtures (**Figure 2e, Figure S9**). This suggests that asymmetries in interaction strengths across the binary mixtures cannot account for the demixing phenotype observed in cells.

Overall, our analyses of the phase diagrams for binary mixtures suggest that the demixing of Whi3-*CLN3* and Whi3-*BNI1* condensates is unlikely to be driven by purely thermodynamic considerations. Therefore, we pivoted to investigating the dynamical aspects of condensate formation in ternary mixtures.

The overall dynamics of phase separation in multicomponent mixtures will be governed by the interplay between timescales associated with molecular transport such as diffusion and the timescales associated with the making and breaking of crosslinks between stickers ^21,22^. These considerations will also be influenced by the stoichiometry of complementary stickers ^23^. Long-lived crosslinks can create long-lived metastable states ^24^. Theory and computations have shown that the totality of these considerations will influence condensate growth and the generation of aspherical morphologies of protein-RNA condensates ^21–26^. We reasoned that colocalization of macromolecular components in condensates requires synchronization of the two relevant timescales. Conversely, asynchronies, caused either by long-lived crosslinks impacting molecular transport ^21^ or significant asymmetries in molecular mobilities ^27^ should give rise to dynamically-controlled phase separation. In these scenarios, the details of how the systems are prepared will determine the likelihoods of mixing versus demixing of macromolecules within condensates.

To test whether dynamical factors lead to the demixing of Whi3-RNA condensates, we analyzed the extents of colocalization within *in vitro* condensates formed in ternary systems comprising two types of RNAs and the Whi3 protein. The concentrations of Whi3 (5 μM) and different RNA molecules (5 nM) were chosen because these are inside the two-phase regimes in binary mixtures (**Figure 2c**). The condensates were formed using two different methods. The first method, termed *delayed*, which replicates the work of Langdon et al.,^7^ involves mixing Whi3 with one of its cognate RNAs, waiting for four hours, and then adding in the second RNA (**Figure 3a**). The second method, termed *simultaneous*, involved adding the two RNAs simultaneously to Whi3 (**Figure 3b**).

**Figure 3:**
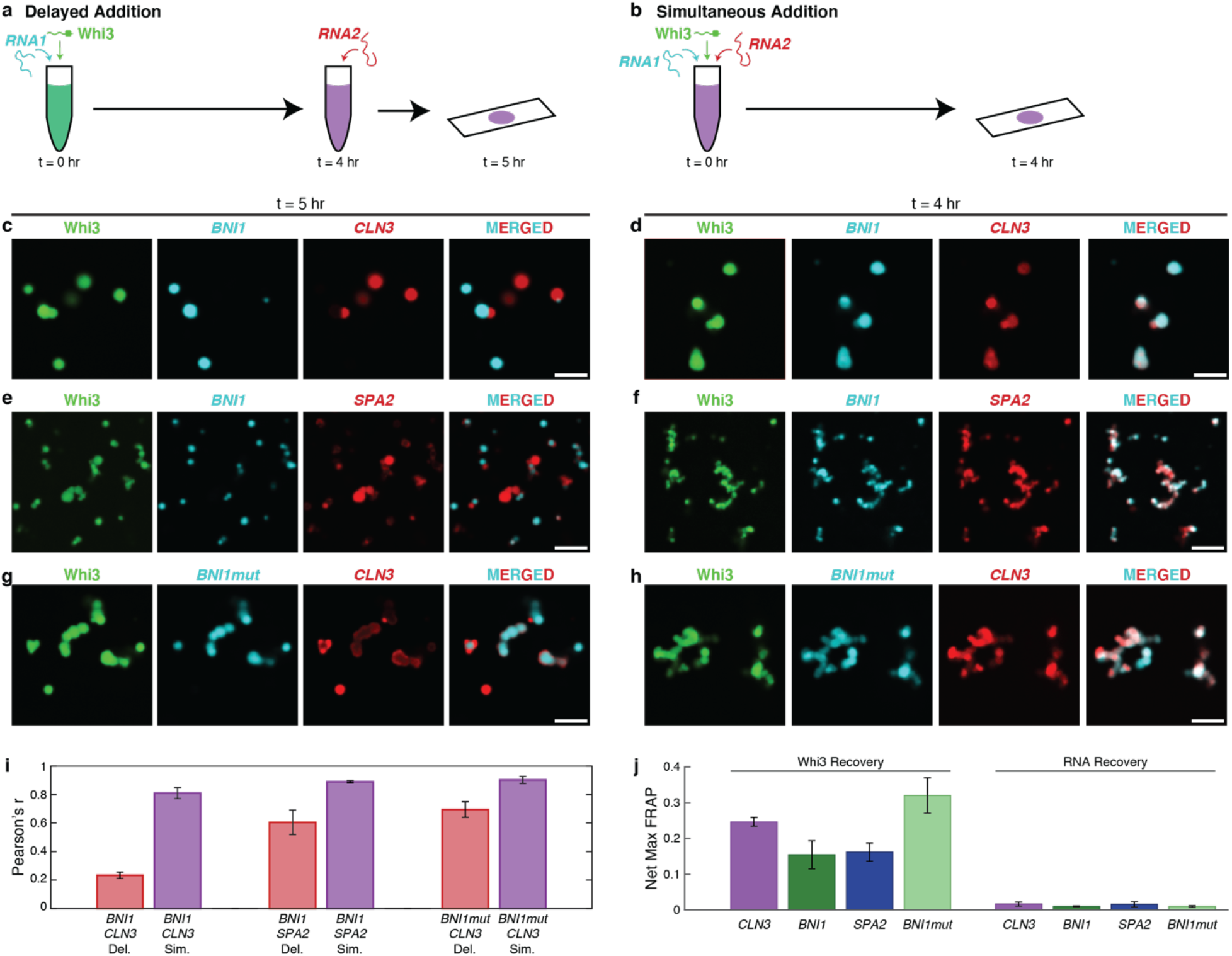
Demixing of condensates in ternary systems is enabled by dynamical arrest caused by metastable traps in binary systems. (a,b) Schematics of the mixing methods of Whi3 with two of its cognate RNAs. Whi3 and RNA concentrations are 5 μM and 5 nM, respectively. (c,d) Confocal images of condensates formed by Whi3, *BNI1*, and *CLN3* in (c) *delayed* versus (d) *simultaneous* modes (e,f) Confocal images of condensates formed by Whi3, *BNI1*, and *SPA2* in (e) *delayed* versus (f) *simultaneous* modes. (g,h) Confocal images of condensates formed by Whi3, *BNI1mut*, and *CLN3* in (g) *delayed* versus (h) *simultaneous* addition. (i) Pearson *r*-values quantifying the colocalization of pairs of RNA molecules. Data for the *delayed* mode are shown in red, whereas data for the *simultaneous* mode are shown in purple. Error bars denote standard errors of mean. (j) Net maximal recovery in normalized fluorescence values from FRAP traces for Whi3 and RNA in binary Whi3-RNA systems. Error bars denote standard error of mean of the Pearson *r* values computed across three replicates.

The extent of colocalization of *CLN3* to pre-formed Whi3-*BNI1* condensates was low when condensates in ternary mixtures were prepared using the *delayed* approach (**Figure 3c,i**). In contrast, *simultaneous* mixing led to a higher degree of colocalization of *CLN3* and *BNI1* in condensates with Whi3 (**Figure 3d,i**). We obtained similar results for ternary mixtures of Whi3, *BNI1*, and *SPA2* though high colocalization is seen even in the delayed scenario, consistent with their tendency to co-localize in cells (**Figure 3e,f,i**). Overall, the extents of colocalization of RNA molecules was highest in the *simultaneous* mode of condensate preparation. These observations suggest that metastable traps form in binary mixtures and these traps hinder the mixing with and colocalization of the ternary component. In addition to differences in colocalization of RNA molecules observed using the *delayed* versus *simultaneous* modes of condensate preparation, we found that the extents of colocalization of *CLN3* and *BNI1* are lower when compared to the extents of colocalization of *BNI1* and *SPA2* ^7^. This is true of either mode of condensate formation and consistent with previous work ^7^.

We also quantified the extents of colocalization achieved in ternary mixtures of Whi3, *CLN3*, and *BNI1mut*. Abrogation of the cognate binding sites for Whi3, increased the extent of colocalization in the *delayed* mode (**Figure 3g,h,i**). This result suggests that cognate binding sites contribute more significantly to long-lived crosslinks when compared to non-cognate sites. As a stringent test of the hypothesis that dynamical influences might be generic drivers of demixing via long-lived metastable states, we studied the extent of colocalization / demixing in a binary mixture that is designed to mimic a ternary system. Specifically, we labeled *CLN3* molecules with two different dyes. In the *simultaneous* mode, we observed a high degree of colocalization between the *CLN3* molecules labeled with different dyes (**Figure S10d, i**). In contrast, in the *delayed* mode, we observed significant diminution of colocalization (**Figure S10c, i**). Similar results were observed for binary mimics of ternary mixtures involving Whi3 and *BNI1* as well as Whi3 and *BNI1*mut (**Figure S10e-i**).

Next, we probed the internal molecular dynamics of Whi3 and different RNA molecules in condensates formed by four different binary mixtures. For this, we measured fluorescence recovery after photobleaching (FRAP) of the labeled components in condensates formed by binary mixtures of Whi3 and RNA molecules in aqueous buffers. In all mixtures, photobleached Whi3 shows partial recovery of its fluorescence, suggesting the presence of mobile and immobile fractions (**Figure 3j, Figure S11a-d**). The fraction of mobile proteins, which corresponds to the maximum recovery, is highest in Whi3-*BNI1mut* condensates, implying that the presence of cognate-binding sites decreases Whi3 mobility. In contrast, RNA molecules are essentially immobile showing undetectable recovery of fluorescence after photobleaching in all the condensates formed by binary mixtures (**Figure 3j, Figure S11e-h**).

FRAP measurements show that Whi3 has slightly more enhanced mobility in the presence of *CLN3* when compared to *BNI1* and *SPA2*. This is true even though Whi3-*CLN3* systems show the greatest degree of demixing upon delayed addition. While these observations are not inherently contradictory, it suggests that there are features other than binary interactions between Whi3 and the RNA molecules that contribute to the extent of mixing / demixing. For example, Langdon et al., proposed that the accessibility of complementary sites among RNA molecules might contribute to demixing ^7^. Indeed, we find that *CLN3* has the smallest number of complementary sites with itself, followed by *BNI1mut* and *BNI1* (**Figure S12**). This suggests that *CLN3* has limited ability to be recruited to preexisting condensates based on RNA-RNA interactions. Additionally, while *BNI1* and *BNI1mut* have a similar number of complementary sites, *BNI1mut* shows a lesser extent of demixing – a result that highlights the contribution of cognate Whi3-binding sites to dynamical demixing.

The overall picture that emerges suggests a push versus pull between two distinct contributions that lead to dynamically controlled demixing of Whi3-RNA condensates. The push toward demixing appears to be caused by long-lived crosslinks between Whi3 and cognate sites on RNA molecules. This competes with a pull away from demixing that is driven by the entropy of mixing that arises from the combinatorics of interactions due to the high valence of non-cognate Whi3-binding sites in *BNI1* and *SPA2*, and the presence of complementary RNA-RNA interaction sites in these molecules.

Finally, we asked if the demixing of Whi3-*CLN3* and Whi3-*BNI1* condensates in live cells of *A. gossypii* is likely to be under dynamical control as predicted from the *in vitro* assays. For this, we engineered an *A. gossypii CLN3^+^/BNI1^+^* strain co-expressing the two transcripts under a common promoter. Co-expression mimics the *simultaneous* mode that we deployed *in vitro* and should enhance colocalization of the two transcripts if the timing of expression influences the extent of co-localization. A bidirectional promoter derived from the *A. gossypii* H2A/B histone locus was chosen to ensure robust co-expression in a wild-type background.

Consistent with previous results ^7^, wildtype expression showed a lack of colocalization between *CLN3* and *BNI1* (**Figure 4a, b**). In contrast, when *CLN3* and *BNI1* were expressed from the same promoter, the extent of colocalization of the two RNAs increased significantly (**Figure 4a,b**). Pixel shift controls suggest that this increase is not solely due to a high density of *CLN3* and *BNI1* in the *CLN3^+^/BNI1^+^* co-expression strain (**Figure S13a**). We also observed a spatial component to the patterns of demixing versus colocalization. Colocalized transcripts are more prominent near the nuclei, which is where the transcripts are synthesized (**Figure S13b**). This suggests that the interplay between spontaneous mixing and dynamically controlled demixing will favor demixed condensates away from the nucleus, whereby longer-lasting RNAs located farther from their production sites are more likely to condense asynchronously with Whi3 thus giving rise to dynamically controlled demixed condensates.

**Figure 4:**
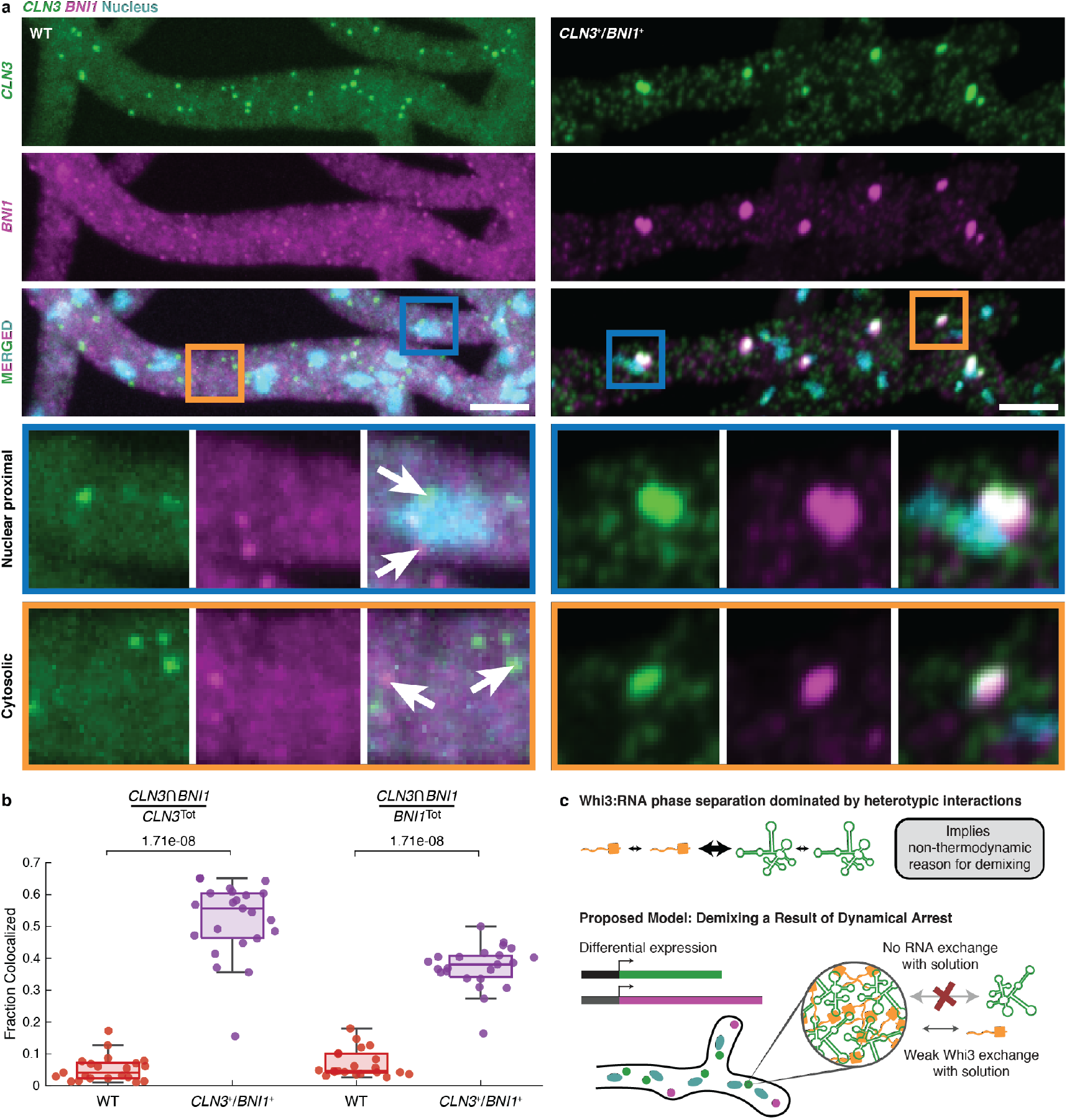
Co-overexpression of *CLN3* and *BNI1* show increase in colocalization. (a) Representative images showing two-color smFISH of *CLN3* (green) and *BNI1* (magenta) in wildtype (WT, left) and CLN3^+^/BNI1^+^ (right) cells. Hoechst signals are shown in cyan, and the scale bar represents 5 μm. Insets show examples of nuclear proximal and cytosolic smFISH signals in the two strains. Arrows highlight individual *CLN3* and *BNI1* spots. (b) Quantification of the colocalizing fractions of *CLN3* and *BNI1* in WT (red) and CLN3^+^/BNI1^+^(purple) cells. (c) Summary schematic of the interactions and arrested dynamics observed for Whi3-RNA systems.

Overall, the results show that enhanced colocalization of *CLN3* and *BNI1* in condensates with Whi3 can be realized by synchronizing their expression *in vivo*. This mimics the *simultaneous* mode of adding the three components *in vitro*. However, in wild-type cells, the expression levels, the locations, and the timings are all different. Therefore, we propose that these dynamical factors rather than purely thermodynamic driving forces, are the primary determinants of demixing of Whi3-*BNI1* and Whi3-*CLN3* condensates in *A. gossypii* (**Figure 4c**).

There is growing interest in the determinants of compositionally distinct coexisting condensates ^6,7,28^, and the driving forces for forming multiphasic, multilayered condensates ^13,29,30^. Proposals based on purely thermodynamic considerations focus on relative interfacial tensions, and hence relative solubilities, as the main determinants of condensate demixing or spatially organized structures of condensates ^20,29–35^. Sequence-intrinsic ^7,20,36,37^ interactions, emulsification ^38^, chemical reactions ^39^, and elastic networks within cells ^40,41^ have also been proposed as regulators of the sizes and compositional specificities of coexisting ribonucleoprotein condensates. Our work adds sequence-intrinsic non-equilibrium dynamical complexities of ribonucleoprotein mixtures to the list of possible controllers of condensate sizes and compositional specificities. Indeed, dynamical control cannot be ruled out in the observed “unblending” of transcriptional condensates that are influenced by repeat expansions that code for intrinsically disordered regions of transcription factors ^28^.

Our findings bolster theoretical predictions ^21,26^ while also highlighting the relevance of viscoelastic phase separation ^27,42^ as a likely mechanism for phase separation *in vitro* and in live cells. In this process, differential mobilities of different macromolecules and timescales associated with competing processes *viz*., molecular transport (influenced by bulk and macromolecule-specific intrinsic viscosities) versus the making and breaking of interactions (the contribute to elasticity) can engender phase separation on different length- and timescales. The large cytoplasm of the filamentous fungus *A. gossyppi*, and quite likely other mammalian cells such as axons ^43^, are likely candidates to leverage dynamical control to enable the formation of coexisting, demixed ribonucleoprotein condensates comprising the same protein and different RNA molecules.

## Acknowledgments

We thank Nadia Erkamp, Mrityunjoy Kar, Tanja Mittag, and members of the Pappu and Gladfelter labs for insightful feedback and critical reading of the manuscript. This work was funded by the Air Force Office of Scientific Research grant (FA9550-20-1-0241 to ASG and RVP), the St. Jude Research Collaborative on the Biology and Biophysics of RNP granules (to RVP), and the National Institutes of Health (F32GM146418-01A1 to MRK).

## Additional information

**Figures S1-S13** are available in the supplementary materials. Details of the computational methods, *in vitro* experiments, cellular measurements, constructs used and the analysis of computed as well measured data are presented in the supplementary material.

**Correspondence and requests for materials** should be addressed to A.S.G. and R.V.P.

