## Supplemental Materials for "Dynamical control enables the formation of demixed biomolecular condensates"

### Methods

### LaSSI Simulations

To investigate minimal models for associative polymers that can lead to equilibrium demixed condensates, we used a customized version of the lattice simulation engine LaSSI [1]. Furthermore, to generate plots and figures we used the common Python packages NumPy, Matplotlib and Seaborn. Adobe Illustrator is used to make the final figures.

### Simulations of ternary systems

For investigating the emergence of demixed condensates given two different RNA species with Whi3 protein, we simulated three different polymer species. The Whi3 protein is modeled as a polymer containing three beads, $W_3$, while the two RNA species are modeled as eight bead polymers, $A_8$ and $B_8$, for *RNA1* and *RNA2* respectively. The beads are connected via implicit linkers with a length of two lattice sites. Using a lattice size of $L = 100$, we placed 5000 $W_3$, 1000 $A_8$, and 1000 $B_8$ molecules within the simulation boxes.

### Interaction Models

To study scenarios that give rise to thermodynamically controlled demixing of condensates generated by three-component systems that mimic the Whi3 protein + *RNA1* + *RNA2* systems, we used multiple interaction models to assess how different interactions among the components affect the overall phase behavior.

#### *Base Case*

To set baseline expectations regarding the phase behaviors of three-component systems containing Whi3 protein and two RNA species, we included heterotypic interactions between the Whi3 protein and the RNAs. This is referred to as the *Base Case*. Stickers interact with pairwise-exclusive interactions, that are equivalent to the binding between the stickers. Given $\epsilon = -2k_BT$ as the energy scale, the interactions between the different stickers for the *Base Case* are defined as

$$\hat{\mathcal{E}}_{Base} = \begin{pmatrix} 0 & \epsilon & \epsilon \\ \epsilon & 0 & 0 \\ \epsilon & 0 & 0 \end{pmatrix}.$$

Here, the $\epsilon_{1,j}$ represents the interactions of the Whi3 protein sticker, $\epsilon_{2,j}$ and $\epsilon_{3,j}$ represent the interactions for *RNA1* and *RNA2*, respectively. Lastly, with this notation, $\epsilon_{i,i}$ represents homotypic interactions, and $\epsilon_{i,i\neq j}$ heterotypic interactions.

*Scaling of homotypic interactions*

To test the effects of homotypic interactions between the RNAs, we added a scaling factor, $h$, and included homotypic interactions among the RNA stickers:

$$\hat{\mathcal{E}}_{Hom.\ Scaling} = \begin{pmatrix} 0 & \epsilon & \epsilon \\ \epsilon & h\,\epsilon & 0 \\ \epsilon & 0 & h\,\epsilon \end{pmatrix},$$

where the scaling factor $h \in \left\{0, \frac{1}{2}, 1, \frac{3}{2}, 2\right\}$ determines the strength of the homotypic interaction for each RNA species with itself. The case where $h = 0$ represents the *Base Case* mentioned above where no interactions occur between the RNAs.

*Asymmetric heterotypic interactions*

To test whether asymmetric interactions between the Whi3 protein and an RNA species could lead to demixing, we applied a scaling factor, $a$, to the heterotypic interactions in the *Base Case*:

$$\hat{\mathcal{E}}_{Asymm} = \begin{pmatrix} 0 & a\,\epsilon & \epsilon \\ a\,\epsilon & 0 & 0 \\ \epsilon & 0 & 0 \end{pmatrix},$$

where the scaling factor $a \in \left\{\frac{1}{2}, 1, \frac{3}{2}, 2, 3\right\}$ determines the strength of the heterotypic interaction between Whi3 protein and *RNA1*. The case where $a = 1$ represents the *Base Case* where there are no differences between RNA species.

*Unphysical repulsion*

To set expectations about the phase behavior of a model that does lead to demixing, we added an additional isotropic repulsive interaction between the RNA species. The isotropic repulsion is a contact potential with a radius of one lattice site or $\sqrt{3}$ lattice units. Given $\epsilon_{rep} = 0.04k_BT \approx \frac{1}{26}k_BT$, we have

$$\hat{\mathcal{E}}_{Base} = \begin{pmatrix} 0 & \epsilon & \epsilon \\ \epsilon & 0 & 0 \\ \epsilon & 0 & 0 \end{pmatrix} \text{ and } \hat{\mathcal{E}}_{rep}^{ISO} = \begin{pmatrix} 0 & 0 & 0 \\ 0 & 0 & s\,\epsilon_{rep} \\ 0 & s\,\epsilon_{rep} & 0 \end{pmatrix},$$

where the scaling factor $s \in \{1,2,5,10\}$ determines the additional heterotypic isotropic repulsion between the different RNA species. Repulsion is disallowed. Here, the RNAs do not repel their own species, but repel the other RNA species, which makes it less likely for different RNA species to be proximal. Again, if $s = 0$, we recover the *Base Case*.

*Simulation protocol*

All simulations start with random initial conditions. For $t = 5 \times 10^7$ MC steps, the simulation temperature is set to $T_{EQ} = 10\,T^*$. A constraining potential is applied to the system which pushes all molecules towards the center of the simulation box. This potential has the form: $V(r,T) = T\,H(r - R_B)\,r^2$, where $r$ is the distance of a given bead from the center of the lattice, $R_B = 35$ lattice units, and $H(r)$ is the Heaviside function. Here, $T = T_{EQ}$ is a set constant. This potential resembles an *indent* style potential as implemented in popular packages such as LAMMPS [2]. All anisotropic/binding interactions are also turned off during this phase of the simulation. After $t_{EQ}$ MC steps, anisotropic interactions are turned on and the temperature is exponentially annealed using an annealing protocol $T(t) = T_0 + T_{EQ}e^{-4\frac{t}{t_{EQ}}}$, to the target temperature of $T_0 = T^*$. As the temperature decreases and $T(t) - T_0$ gets lower than a threshold of $0.005$, the temperature is set to $T_0$ and the biasing potential is turned off. The simulations are run for $t = 1 \times 10^{10}$ MC steps, and samples are only taken in the last half of each run. Samples are taken every $f_{data} = 2.5 \times 10^6$ MC steps, which result in $2000$ samples for each simulation temperature. Three replicates per condition were used. The standard error of the mean between replicates is used as a measure of uncertainty.

**Table S1: Frequencies of Monte Carlo moves for three-component mixture simulations**

| Move Type | Normalized By Min | Frequency |
|---|---|---|
| Rotation | 500 | 4.75 |
| Local | 5000 | 47.52 |
| Co-Local | 1000 | 9.50 |
| Multi-Local | 500 | 4.75 |
| Chain Reptation | 500 | 4.75 |
| Chain Translation | 1000 | 9.50 |
| Aniso. Cluster Translation (Small) | 10 | 0.10 |
| Aniso. Cluster Translation (Large) | 1 | 0.01 |
| Chain Pivot | 1000 | 9.50 |
| Double Pivot | 1000 | 9.50 |
| Cluster Translation (Small) | 10 | 0.10 |
| Cluster Translation (Large) | 1 | 0.01 |

Details about the different moves can be found in the original LaSSI work, Choi et. al. [1], and the Supporting Information Appendix of Kar et. al. [3].

#### Generation of two-component phase diagrams with LaSSI

For generating the two-component phase boundaries, we sampled a large set of concentrations and stoichiometries and explicitly measure the coexisting densities, when condensates are formed in the simulations. Since we have two explicit components, the system composition is determined by the numbers of each molecule, and the overall concentration of the system. We fixed the total number of molecules to 5000 and changed the stoichiometry by changing the ratio between the two components. This results in 22 different stoichiometries. The simulation box size is then used to set the total concentration of the system. For each pair of molecule numbers, we have five different box-sizes. Lastly, for each pair of molecule numbers, $(n_1, n_2)$, and box-size $L$, we also sample the $(n_2, n_1)$ composition. This gives us a total of 220 independent compositions for a given interaction model.

**Table S2: Numbers of molecules and box-sizes for two-component phase diagrams**

| **Molecule 1** | **Molecule 2** | $L_1$ | $L_2$ | $L_3$ | $L_4$ | $L_5$ |
|---|---|---|---|---|---|---|
| 185 | 4815 | 46 | 55 | 67 | 81 | 99 |
| 256 | 4744 | 51 | 61 | 74 | 90 | 109 |
| 442 | 4558 | 61 | 73 | 89 | 108 | 131 |
| 638 | 4362 | 69 | 83 | 101 | 122 | 148 |
| 834 | 4166 | 76 | 92 | 111 | 135 | 163 |
| 1030 | 3970 | 81 | 98 | 118 | 144 | 174 |
| 1303 | 3697 | 88 | 106 | 129 | 156 | 189 |
| 1618 | 3382 | 95 | 115 | 139 | 168 | 204 |
| 1814 | 3186 | 98 | 118 | 143 | 174 | 211 |
| 2206 | 2794 | 105 | 127 | 154 | 186 | 226 |
| 2500 | 2500 | 109 | 132 | 159 | 193 | 234 |
| 2794 | 2206 | 114 | 138 | 167 | 202 | 245 |
| 3186 | 1814 | 119 | 144 | 174 | 211 | 256 |
| 3382 | 1618 | 121 | 146 | 177 | 215 | 260 |
| 3697 | 1303 | 125 | 151 | 183 | 222 | 269 |
| 3970 | 1030 | 128 | 155 | 187 | 227 | 275 |
| 4166 | 834 | 130 | 157 | 190 | 231 | 280 |
| 4362 | 638 | 132 | 159 | 193 | 234 | 284 |
| 4558 | 442 | 134 | 162 | 196 | 238 | 288 |
| 4744 | 256 | 136 | 164 | 199 | 241 | 293 |
| 4904 | 96 | 137 | 165 | 201 | 243 | 295 |
| 4950 | 50 | 138 | 167 | 202 | 245 | 297 |

*Simulation protocol for two-component systems*

All simulations start with random initial conditions. For $t_{EQ} = 5 \times 10^7$ MC steps, the simulation temperature is set to $T_{EQ} = 100\,T^*$. A constraining potential is applied to the system which pushes all molecules towards the center of the simulation box. This potential has the form: $V(r,T) = \Delta T \cdot r^2$, where $r$ is the distance of a given bead from the center of the lattice, $\Delta T$ is the temperature difference between the current simulation temperature and $T_0 = T^*$, the first target temperature. All anisotropic/binding interactions are turned off during these initial steps. After $t_{EQ}$ MC steps, anisotropic interactions are turned on and the temperature is exponentially annealed as such $T(t) = T_0 + T_{EQ} e^{-4\frac{t}{t_{EQ}}}$, to the target temperature of $T_0 = T^*$. As the temperature decreases $\Delta T$ gets lower than a threshold of $0.005$, the temperature is set to $T_0$, and the biasing potential is turned off. The simulations are run for $t = 2 \times 10^9$ MC steps, and samples are only taken in the last half of each run. Samples are taken every $f_{data} = 1 \times 10^6$ MC steps, which result in $1000$ samples for each run condition. Two replicates per condition were used.

**Table S3: MC move frequencies for two-component phase diagrams**

| Move Type | Normalized by Min | Frequency |
|---|---|---|
| Rotation | 500 | 14.24 |
| Local | 1000 | 28.48 |
| Co-Local | 666 | 18.99 |
| Multi-Local | 500 | 14.24 |
| Chain Reptation | 333 | 9.49 |
| Chain Pivot | 166 | 4.75 |
| Double Pivot | 166 | 4.75 |
| Chain Translation | 166 | 4.75 |
| Anisotropic Cluster Translation (Small) | 10 | 0.28 |
| Anisotropic Cluster Translation (Large) | 1 | 0.03 |

**Analysis of data from LaSSI simulations**

*Generating radial density distributions*

Given each simulation, we explicitly calculated the radial density profiles of each component from the center-of-mass (COM) of the system, and from the COM of the largest cluster of each component. The density profiles are generated by first computing a number histogram of beads, $H(r_n)$, from a given COM, with bin-width $0.25$, from $r = 0$ to $r = \frac{\sqrt{3}\,L}{2}$, given a lattice size $L$. To normalize this number histogram, we explicitly calculated the number histogram of lattice sites, $H_0(r_n)$ for a given lattice size $L$ with periodic boundaries, and a bin-width of $0.25$. Thus, the density profiles are calculated as:

$$\rho(r_n) = \frac{H(r_n)}{H_0(r_n)}.$$

*Coexisting densities*

To calculate the coexisting densities for the two-component systems, given a component $i$, we used the system COM to calculate the density profiles $\rho_i(r_n)$. We then averaged over the first 13 bins, and 20 bins near the end of the simulation box, avoiding the last 15 bins.

*Measure for demixing in three-component systems*

Let component $i$ be the COM component, and let $j$ be the component for which we are calculating the density, then $\rho_{ij}(r_n)$ denotes the density profile of component $j$, given component $i$ as the COM. Given a density profile $\rho_{ij}(r)$, we can generate a normalized distribution, $\widetilde{\rho_{ij}}(r)$, such that

$$\sum_{n=0}^{N_{bins}} \widetilde{\rho_{ij}}\,(r_n) = 1.$$

Then, using the Hellinger Distance [4], we can define a measure for demixing:

$$\mathcal{D}_{ij} = D_H(\widetilde{\rho_{ii}}\,, \widetilde{\rho_{ij}}),$$

where the Hellinger Distance, $D_H(P, Q)$, given distributions $P(x)$ and $Q(x)$, is defined as

$$D_H = \sqrt{1 - BC(P, Q)},$$

and where $BC$, the Bhattacharyya Coefficient, is defined as

$$BC(P,Q) = \sum_{x \in X} \sqrt{P(x)Q(x)}.$$

Combining all definitions,

$$\mathcal{D}_{ij} = \sqrt{1 - \sum_{n=0}^{N_{bins}} \sqrt{\widetilde{\rho_{ii}}(r_n)\widetilde{\rho_{ij}}(r_n)}}\,.$$

Thus, $\mathcal{D}_{ij}$ acts as a measure for the demixing between components $i$ and $j$.

**Extracting goodness-of-fit from phase diagrams obtained using coarse-grained simulations**

Phase diagrams were plotted in terms of sticker concentration. The dilute phase arm of the phase diagram (**x**, **y**) was then mirrored by also plotting (**y**, **x**). The points that defined the dilute phase arm and the mirrored data ([**x y**], [**y x**]) were then fit to an ellipse using the guaranteed ellipse fitting method of Szpak et al.,[5] and the mean square of the residuals $R^2$ of the fit was extracted.

**Protein purification and tagging**

Full-length Whi3, with a N-terminal 6x His tag and TEV cleavage site, in BL21 cells from NEB in TB media (Terrific Broth) was induced with IPTG to a final concentration of 1mM at an $OD_{600}$ of 0.6 – 0.7 before being expressed overnight at 18˚C. These cells were then lysed in lysis buffer (1.5 M KCl, 50 mM HEPES pH 8.0, 20 mM Imidazole, 5 mM BME) containing 10mg of lysozyme, one Roche cOmplete™ protease inhibitor cocktail tablet, and PMSF at a final concentration of 200 µM. The resulting lysate was sonicated on ice with a Branson SFX550 at 30% strength alternating one second on, two seconds off for one minute. This was repeated five times, swirling the lysate gently between each sonication. The lysate was spun down and the supernatant passed over a HisTRAP FF column (Cytiva) on an ÄKTA pure 25 L (GE). The bound protein was eluted in 150 mM KCl, 50 mM HEPES pH 8.0, 200 mM Imidazole, and 5 mM BME The eluate then cleaved with 200 µg of TEV [pRK793 was a gift from David Waugh (Addgene plasmid # 8827 ; http://n2t.net/addgene:8827 ; RRID:Addgene_8827)]. The cleaved supernatant was concentrated in 3 KDa Amicon® centrifugal filter units (Millipore Sigma), and then injected onto a HiLoad 16/600 Superdex 200pg column (Cytiva). Untagged Whi3 was then dialyzed into storage buffer (200 mM KCl, 50 mM Tris pH 8.0, 5 mM BME) and concentrated using 3 Kda Amicon® centrifugal filter units. For tagged Whi3, Alexa Fluor 488 NHS Ester (ThermoFisher) in DMSO was added at a ratio of 4:1 and incubated at room temperature with continuous mixing for one hour in the absence of light. The tagged Whi3 was then loaded onto a HiLoad 16/600 Superdex 200pg column with storage buffer, and the fractions concentrated with 3 Kda Amicon® centrifugal filter units.

**RNA production, purification, and tagging**

Plasmids in which the T7 promoter sequence was placed upstream of the coding regions for *CLN3*, *BNI1*, and *SPA2* were linearized with restriction enzymes to obtain a linearized template. *CLN3* RNA was transcribed with HiScribe™ T7 Quick High Yield RNA Synthesis Kit (NEB). *BNI1* and *SPA2* RNA were transcribed using Hi-T7® RNA Polymerase and Reaction Buffer (NEB) in lieu of T7 polymerase and buffer. For labeled RNA, 0.1 µL of 5 mM Cy3-UTP or Cy5-UTP (Cytiva) was added to each transcription reaction. After transcription, each reaction was treated with Dnase before being purified with Monarch® RNA Cleanup Kit (NEB).

***In vitro* measurements of phase boundaries**

384-well plates (Ibidi) were passivated for 15 minutes with 0.1% Tween-20 before being rinsed thrice with droplet buffer (150 mM KCl, 50 mM Tris pH 8.0, 5 mM BME). Untagged Whi3 protein and tagged RNA were diluted in droplet buffer and mixed to obtain the desired final concentration of protein and RNA. After incubation at room temperature for one hour, the samples were visualized on a Zeiss Axiovert 200M with a C-Apochromat 40X 1.2NA water objective.

**Extracting parameters from *in vitro* phase diagrams**

First, we extracted the n points along the lower boundary of the two-phase regime $\mathbf{x_P}$=($x_{P1}$ $x_{P2}$ … $x_{Pn}$) and $\mathbf{y_P}$=($y_{P1}$ $y_{P2}$ … $y_{Pn}$), and the m points along the upper boundary of the one-phase regime $\mathbf{x_N}$=($x_{N1}$ $x_{N2}$ … $x_{Nm}$) and $\mathbf{y_N}$=($y_{N1}$ $y_{N2}$ … $y_{Nm}$). Here, the subscript N and P represent points in the one- (**N**o phase separation) and two-phase (**P**hase separation) regimes, respectively. We then assumed the system is dominated by heterotypic interactions. Under this assumption, we can rescale the RNA concentration ($\mathbf{y_P}$ and $\mathbf{y_N}$) by titrating through a scaling factor, *s*, to find the highest degree of symmetry. The best *s* should have the lowest overlap between the area defined by ($\mathbf{x_N}$, $s\mathbf{y_N}$) and ([$\mathbf{x_P}$ $s\mathbf{y_P}$], [$s\mathbf{y_P}$ $\mathbf{x_P}$]). If heterotypic interactions dominate, then there should be minimal overlap between the area defined by the upper boundary of the one-phase regime and the area defined by the lower boundary of the two-phase regime plus its mirrored data. To determine the overlap area, the MATLAB function *boundary* was used to define the boundaries of ($\mathbf{x_N}$, $s\mathbf{y_N}$) and ([$\mathbf{x_P}$ $s\mathbf{y_P}$], [$s\mathbf{y_P}$ $\mathbf{x_P}$]) and the MATLAB function *polyshape* was used to create polygons defined by the boundaries. Then, we extracted the intersection of the two polygons using the MATLAB function *intersect* and calculated the area of this overlap region using the MATLAB function *area*. Then, the apparent valence of the RNA was taken to be the *s* value that yields the minimum overlap area. The boxplots in **Figure 2f** correspond to the range in apparent valence values if we allow for a five percent change in the minimum area. Lastly, for the *s* that corresponds to the minimum overlap we fit an ellipse to ([$\mathbf{x_P}$ $s\mathbf{y_P}$], [$s\mathbf{y_P}$ $\mathbf{x_P}$]) using the guaranteed ellipse fitting method of Szpak et al., [5] and extracted the $R^2$ of the fit.

***In vitro* colocalization imaging**

384-well plates (Cellvis) were passivated for 15 minutes with 0.1 % Tween-20 before being rinsed thrice with droplet buffer (150 mM KCl, 50 mM Tris pH 8.0, 5 mM BME). For each sample, protein and RNA were diluted to 5 µM and 5 nM, respectively in droplet buffer. For simultaneous samples, tagged Whi3 and tagged RNAs were added in quick succession to a low protein binding microcentrifuge tube (ThermoFisher) and allowed to incubate in darkness at room temperature for four hours before being transferred to a passivated well and imaged on a Nikon Eclipse Ti2 equipped with a Yokogawa CSU-X1 Spinning Disk, 60X oil objective using Nikon Type F oil, and a Hamamatsu ORCA-Flash4.0 V3 camera. For delayed samples, tagged Whi3 and one of two tagged RNAs were mixed in a low protein binding microcentrifuge tube and allowed to incubate in darkness at room temperature for four hours. Then the second RNA was added and allowed to incubate for one hour before being transferred to a passivated well and imaged on the system described above.

***In vitro* colocalization analysis**

Images from three random fields of view for each sample were collected and cropped to 40,000 $px^2$ (approximately 469 $\mu m^2$) in Fiji [6] The images were then split into separate channels and analyzed with the Coloc2 plugin to obtain the Pearson correlation coefficient (PCC). We employed a bisection threshold regression and 1,000 randomizations with a PSF of 3.0. The PCC was taken for each of the three fields of view and plotted with standard error.

**Fluorescence recovery after photobleaching**

384-well plates (Cellvis) were passivated for 15 minutes with 0.1 % Tween-20 before being rinsed thrice with droplet buffer (150 mM KCl, 50 mM Tris pH 8.0, 5 mM BME). Either tagged Whi3 or RNA is mixed with untagged RNA or Whi3 respectively and allowed to incubate for one hour at room temperature. Samples were imaged on a Nikon A1Rsi with a 60X oil objective using Cargille Type B oil. Laser strength for photobleaching was 50 % of maximum intensity of the wavelength corresponding to the fluorophore used. Photobleaching occurred for five seconds with recovery measured for three minutes utilizing the laser corresponding to the fluorophore used.

The Region of Interest (ROI) and Time Analysis tools in Nikon Imagine Software (NIS) were used to trace the fluorescence intensity of three regions – the photobleached spot, an unbleached condensate, and a region with no condensates (i.e., background). Intensity changes over time were evaluated for all three regions and only acquisitions wherein the intensity of the unbleached condensate deviated less than 5 % were used for analysis. Raw intensity values of the photobleached spots from three acquisitions were averaged, subtracted from background, and normalized to the maximum average value. The "max" recovery value corresponds to the maximum value among all the recovery values (omitting the pre-bleach value). Standard error of the mean (SEM) was calculated for individual time points across the three acquisitions and for the maximum intensity values across three acquisitions. SEM was calculated by taking the standard deviation of time-point-matched intensity values or maximum intensity values and dividing this value by the square root of three (i.e., the n of acquisitions). FRAP traces were generated in matplotlib.

**Complementary site analysis**

GUUGle (https://bibiserv.cebitec.uni-bielefeld.de/guugle) was utilized to identify complementary sites between pairs of RNAs. Here, exact matches of length 11 or longer were identified using G-C, A-U, and G-U base pairing. Additionally, for the CLN3-BNI1, BNI1-SPA2, CLN3-CLN3, and BNI1-BNI1 pairs the mean SHAPE value was calculated over all nucleotides in the complementary sequences using the data from Langdon et al.,. [7]

**smFISH and microscopy**

RNA smFISH was performed as previously described [8]. Ashbya cells were grown by inoculating dirty spores in 50 mL Ashbya full medium (AFM), for wild type or AFM supplemented with G418 (200 µg/mL) for the strain overexpressing *CLN3* and *BNI1* in a 500 mL baffled flask. Cells were grown shaking at 110 rpm at 30 °C for 16 h. After formaldehyde fixation and ethanol permeabilization, cells were probed with custom FISH probes from Stellaris. TAMRA-labeled probes against agCLN3 and Quasar-670 labeled probes against agBNI1 were both used at a final concentration of 2.5 nM and hybridized simultaneously at 37 ºC overnight. Nuclei were stained with 5 µg/mL Hoechst in Wash Buffer (2x SSC, 10 % v/v deionized formamide). 20 µL of cells were then mounted in 20 µL Vectashield mounting medium (Vector Laboratories, H-1000-10), sealed with a coverslip and imaged.

Imaging was performed on a Nikon Ti-E stand using a Yokogawa CSU-W1 spinning disk confocal unit. Images were acquired on a Plan-Apochromat 60x / 1.49NA oil-immersion objective using a Zyla sCMOS camera (Andor) on Nikon NIS-Elements software v.4.60.

***In vivo CLN3* and *BNI1* colocalization analysis**

*CLN3* and *BNI1* spots were identified using the BIGFish Spot detection algorithm [9] that treats puncta as local maxima in the smFISH channels given a specified object radius. Puncta (spot) detection can be performed in two methods: 1) a single function call that enumerates the spots directly, or 2) a series of intermediate detection steps that yields the same results as in method 1, but with more debugging information. The latter option was chosen as it provided more data for troubleshooting the development of the spot detection pipeline. Using the latter method for detecting spots, a spot radius, $r$, of 150 nm (1.389 pixels) was used for detection. Maximum intensity projections were used, and images with uneven illumination and few regions of non-overlapping hyphae were discarded. For each image in which the number of *CLN3* and *BNI1* spots were within an order of magnitude, the centers of identified RNA spots were collected. Then, a *CLN3* spot was determined to be colocalized with a *BNI1* spot if any *BNI1* center was a distance less than $2r$ away from the *CLN3* center. Fraction colocalized then refers to the number of all *CLN3* spots that had at least one *BNI1* spot less than $2r$ away divided by the total number of *CLN3* spots. For the pixel shift analysis, the centers of *CLN3* spots were shifted by $2r$ in both the x and y direction. Then, using the shifted *CLN3* spot centers we determined if any *BNI1* spot was less than $2r$ away.

Furthermore, nuclear proximity was determined by first identifying nuclei using the Cellpose 2.0 Python package [10]. Specifically, we ascribed a minimum object diameter of 10 voxels or 1.08 μm for the detection of nuclei regions of interest and performed segmentation. Larger object diameters perform poorly and detect false positives. Then, the centroids and areas of the nuclei masks were extracted using the Scikit-Image Python package [11]. Using the collected centroids and areas, we then determined whether the center of a *CLN3* spot was less than $r+R$ away from the center of any nucleus. Here, $R$ is the nucleus radius in pixels and was estimated to be $\sqrt{(A/\pi)}$, where $A$ is the area of the nucleus. *CLN3* spots less than $r+R$ away from the center of any nucleus were defined to be nuclear proximal and *CLN3* spots greater than or equal to $r+R$ where defined to be not nuclear proximal. Using these classifications, we then determined the fraction of *CLN3* spots colocalized with *BNI1* split by nuclear proximal and not nuclear proximal where colocalization was defined as any *BNI1* spot center less than $2r$ away from the *CLN3* center. The same analyses were also performed using *BNI1* as the reference RNA.

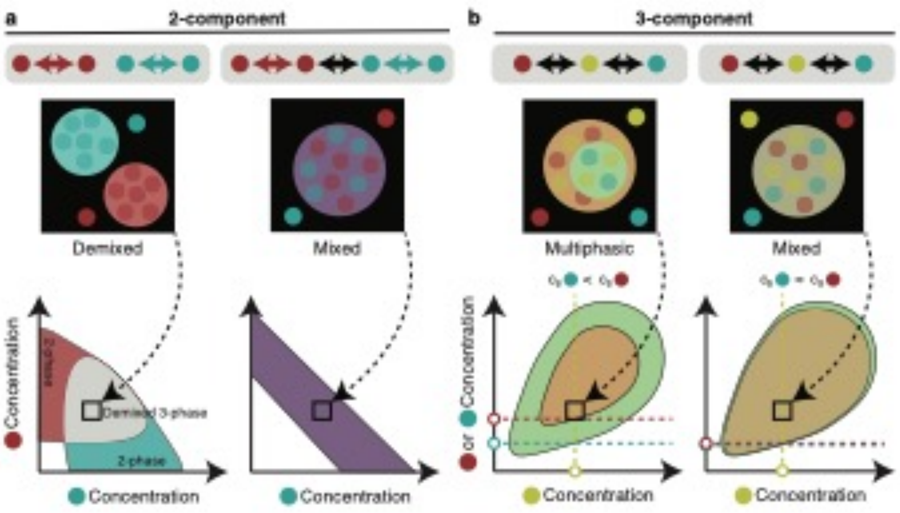


**Figure S1: The interplay between homotypic and heterotypic interactions dictates the thermodynamic preferences for mixed vs. demixed phase behavior.** (a) Left: Example phase behavior for a two-component system in which only homotypic interactions are present. Non-grey color regimes of the phase diagram denote a single condensate type is formed at these conditions, whereas the grey region denotes the region where demixed condensates can form. Right: Example phase behavior of a two-component system in which homotypic and heterotypic interactions are equivalent. Black arrows denote heterotypic interactions, whereas colored arrows denote homotypic interactions. (b) Example phase behavior of a three-component system in which the interaction strength of the yellow component with the two other components is asymmetric (left) or symmetric (right). Green area is the two-phase regime of the two-component system of the yellow and blue molecules, whereas the brown-orange area is the two-phase regime of the two-component system of yellow and red molecules. Here, $c_s$ denotes the saturation concentration at the given starting concentration of the yellow molecule. Drawn to summarize the results of Lu et al., [12].

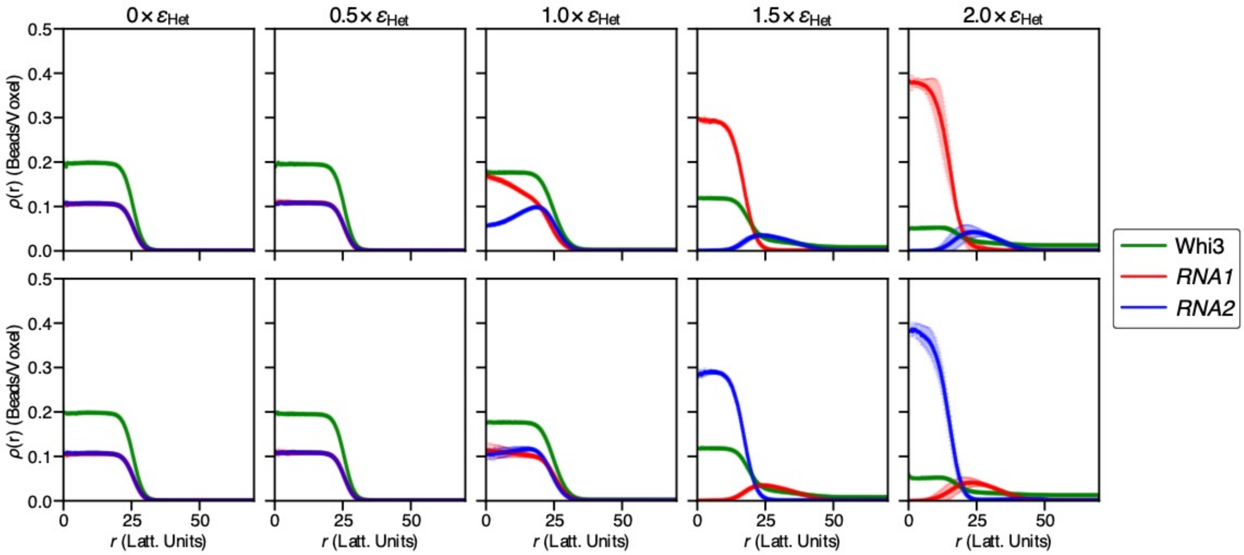


**Figure S2: Density profiles from LaSSI lattice-based simulations of a *RNA1* (red), *RNA2* (blue), Whi3 (green) system in which the strength of homotypic interactions of *RNA1* and *RNA2* are titrated as a function of the Whi3-*RNA* interaction strength.** Here, $\varepsilon_{\text{Het}} = -2k_BT$ and refers to the interaction strength between Whi3-*RNA1* and Whi3-*RNA2*. Each column denotes the homotypic *RNA1*-*RNA1* and *RNA2*-*RNA2* interaction strength as a function of $\varepsilon_{\text{Het}}$. The top and bottom rows show the density profiles from the reference of *RNA1*'s and *RNA2*'s center-of-mass, respectively.

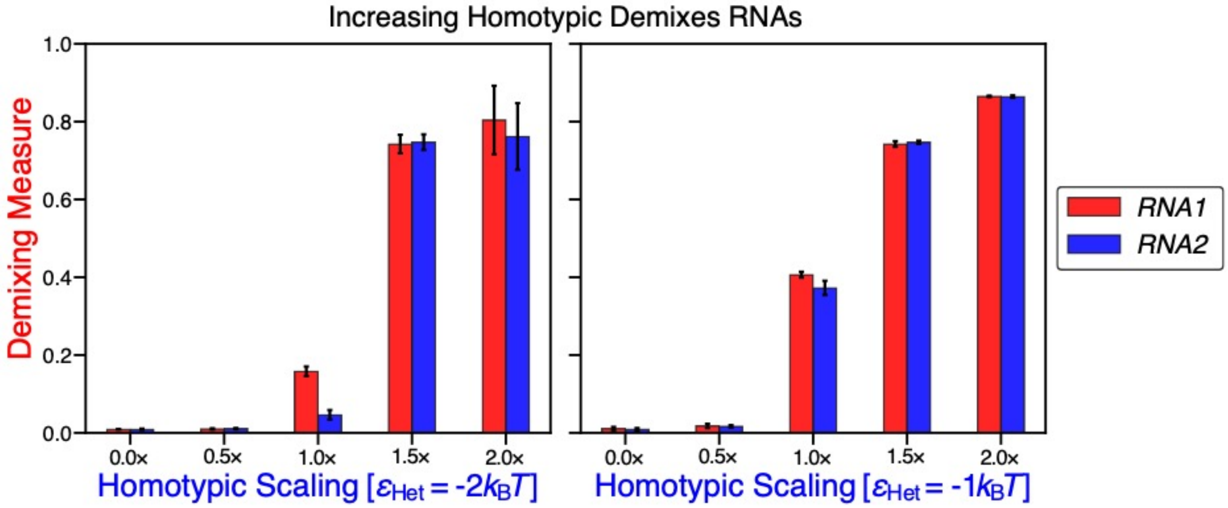


**Figure S3: Demixing measure as a function of homotypic interactions of *RNA1* and *RNA2*.**

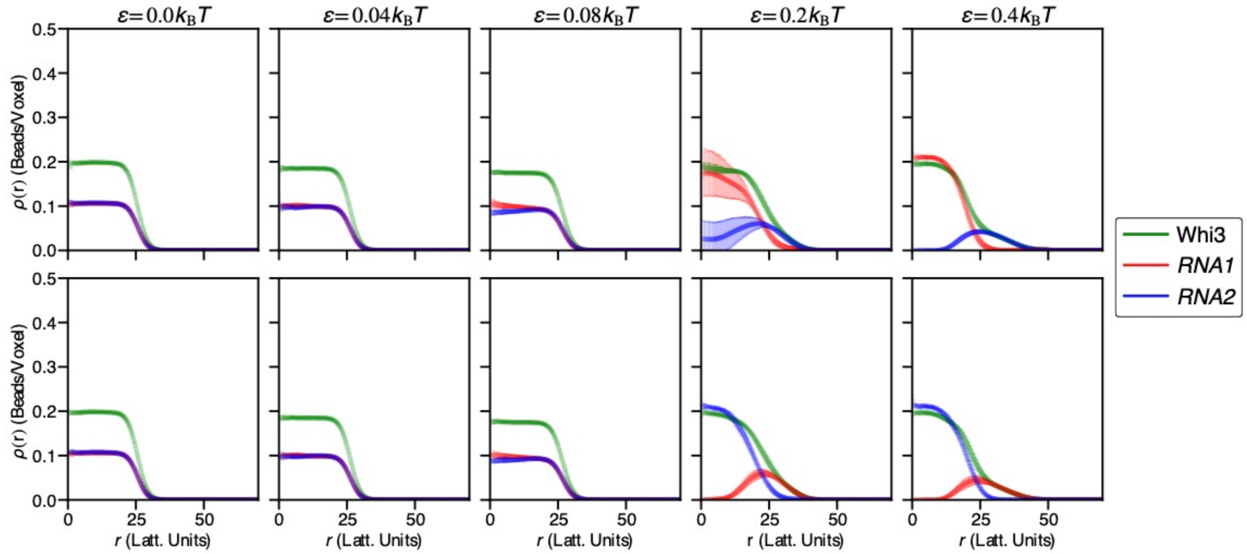


**Figure S4: Density profiles from LaSSI lattice-based simulations of a *RNA1* (red), *RNA2* (blue), Whi3 (green) system in which the strength repulsion between *RNA1* and *RNA2* is titrated.** Here, each column denotes the heterotypic *RNA1*-*RNA2* repulsion strength denoted by ε. The Whi3-*RNA* heterotypic interaction strength is set to $\varepsilon_{\text{Het}}$ = -2$k_BT$. The top and bottom rows show the density profiles from the centers-of-mass *RNA1* and *RNA2*, respectively.

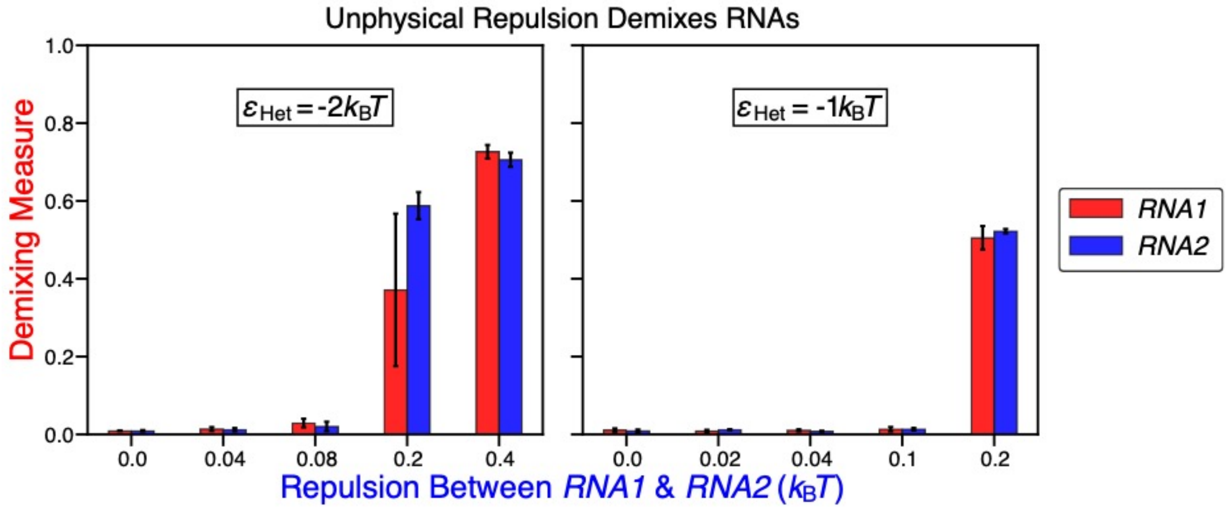


**Figure S5: Demixing measure as a function of the repulsion strength between *RNA1* and *RNA2*.**

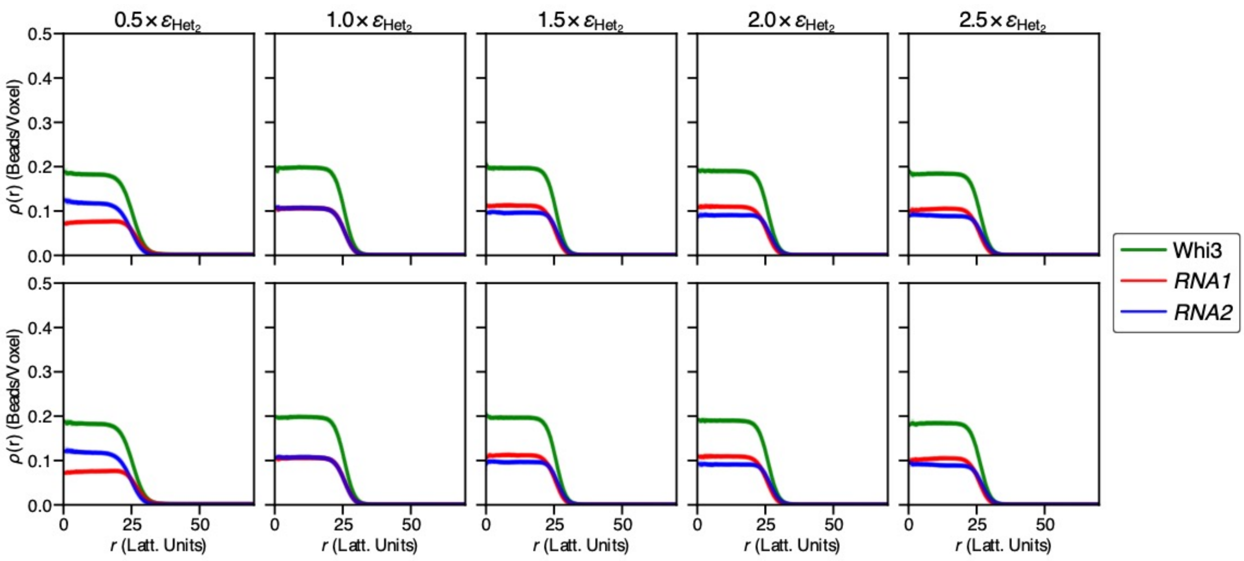


**Figure S6: Density profiles from LaSSI lattice-based simulations of a *RNA1* (red), *RNA2* (blue), Whi3 (green) system in which the strength of the Whi3-*RNA1* interaction is titrated as a function of the Whi3-*RNA2* interaction strength.** Here, $\varepsilon_{\text{Het2}}$ = -2$k_BT$ and refers to the interaction strength between Whi3-*RNA2*. Each column denotes the Whi3-*RNA1* interaction strength as a function of $\varepsilon_{\text{Het2}}$. The top and bottom rows show the density profiles from the centers-of-mass of *RNA1* and *RNA2*, respectively.

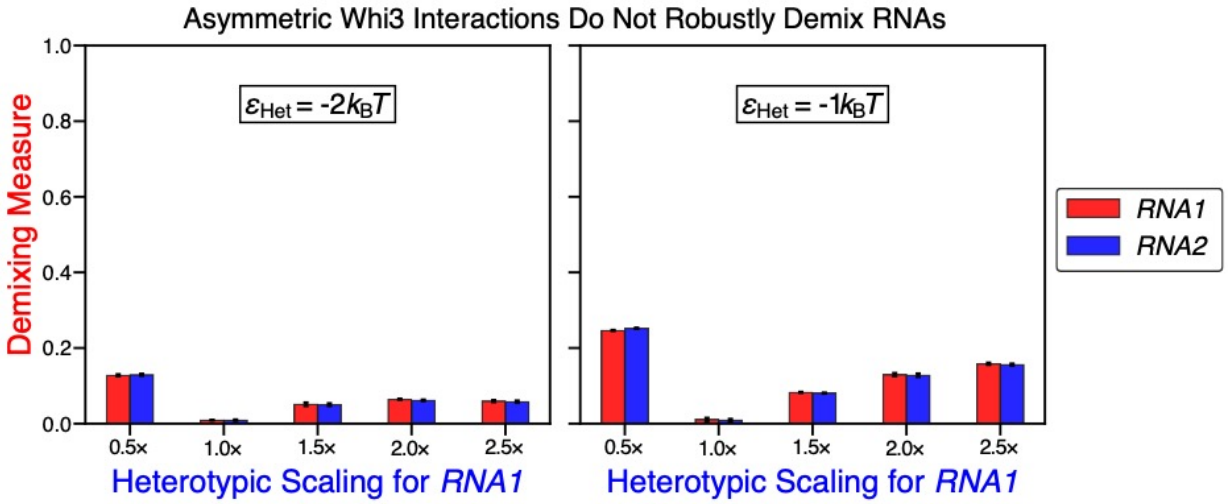


**Figure S7: Demixing measure as a function the heterotypic interaction strength of Whi3-*RNA2*, ε.**

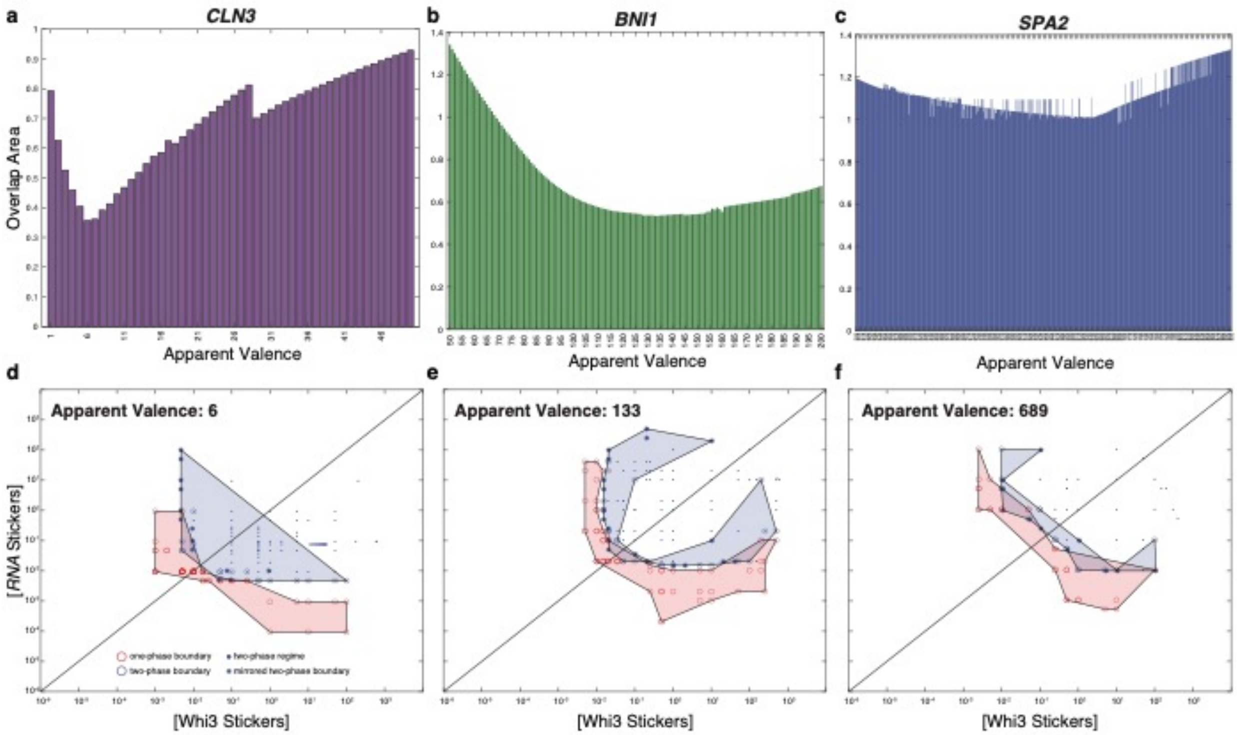


**Figure S8: Dilute and dense phase overlap titration analysis.** (a-c) Overlap area of the one-phase and two-phase boundary regimes given the apparent valence of the *CLN3*, *BNI1*, and *SPA2*, respectively (see Methods). (d-f) Plot of the one-phase and two-phase boundary regimes for the apparent valence with the lowest overlap area. One-phase regimes and the two-phase boundary areas are shown in red and blue, respectively.

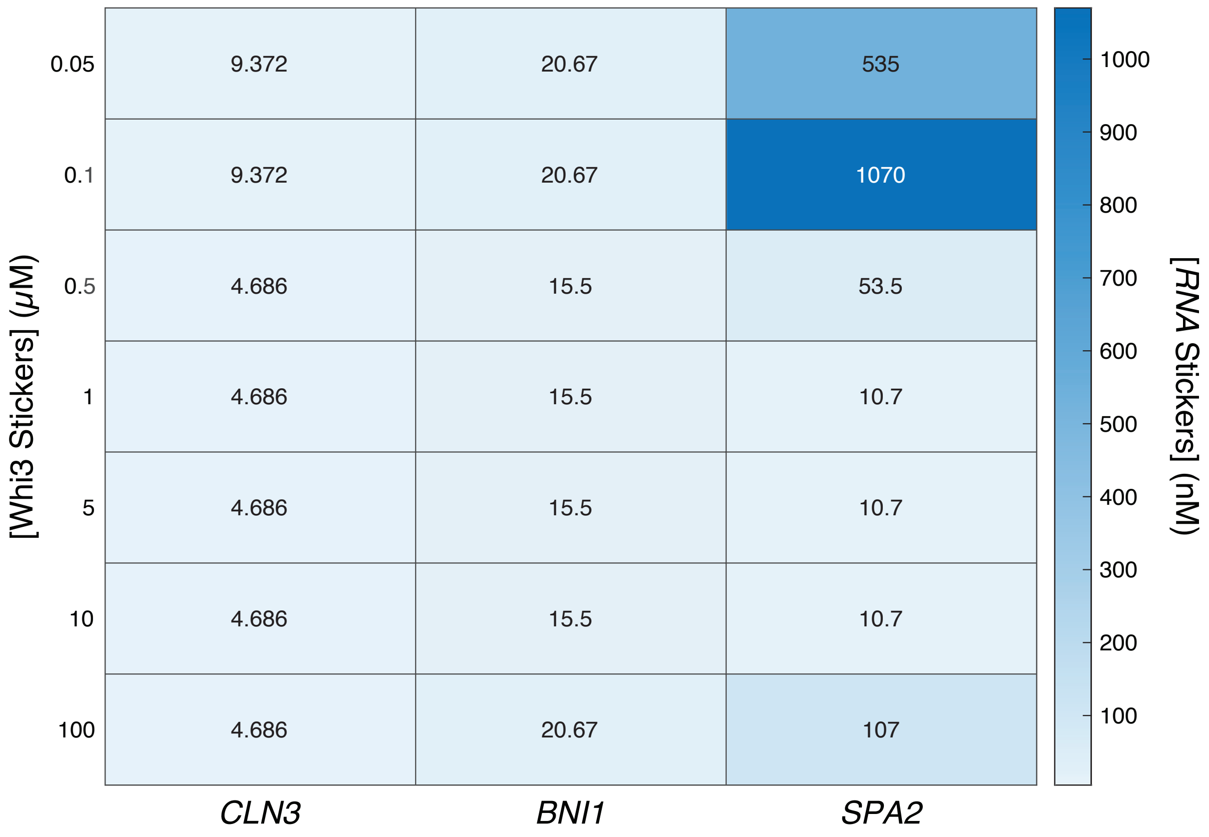


**Figure S9: Dilute arm concentration phase boundary for Whi3 with *CLN3*, *BNI1*, and *SPA2*.**

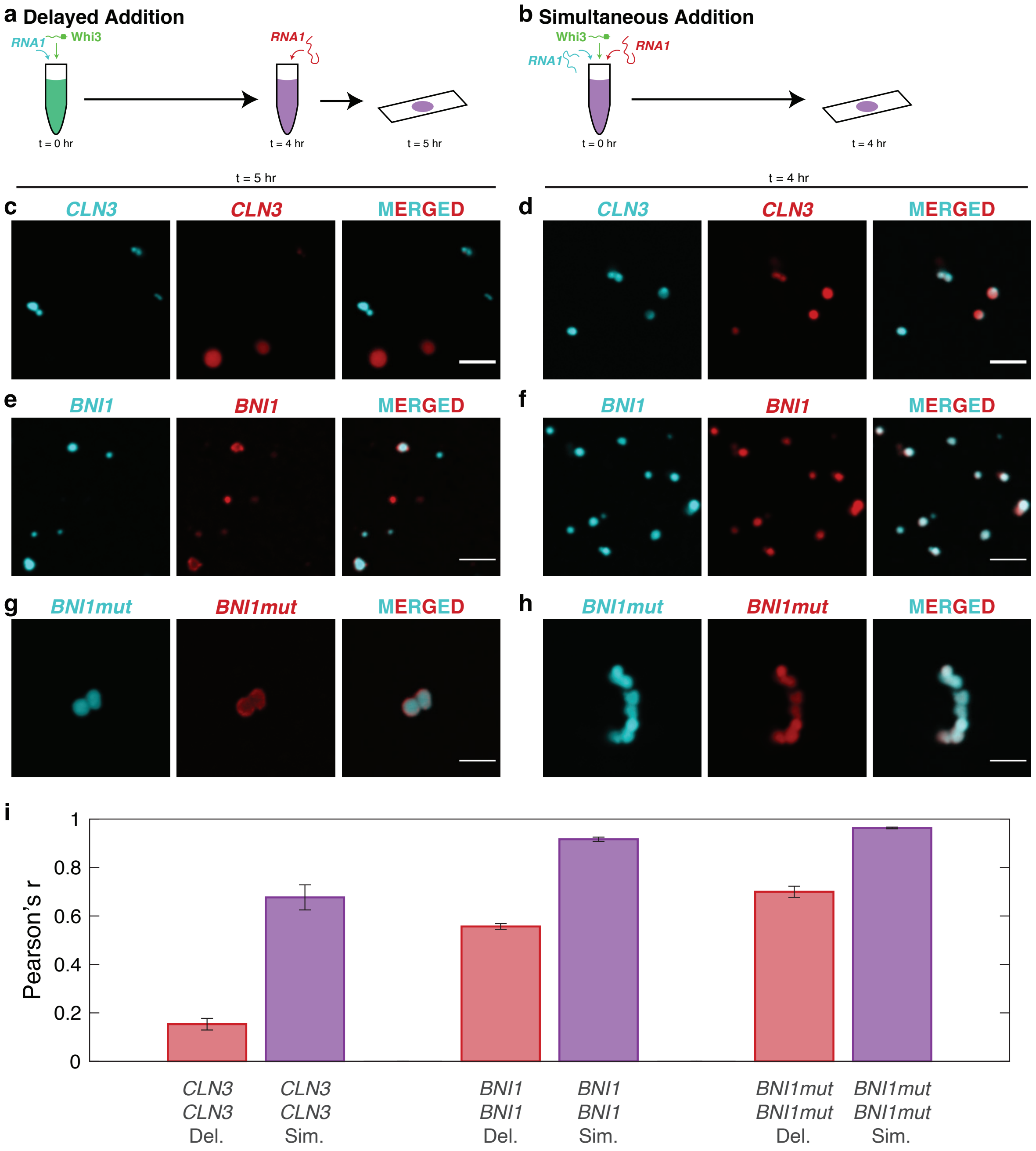


**Figure S10: *In vitro* colocalization of the same RNA in two component Whi3-RNA systems.** (a,b) Schematics of the mixing methods of Whi3 with one of its cognate RNAs labeled with different dyes. Whi3 and RNA concentrations are 5 µM and 5 nM, respectively. (c,d) Confocal images of Whi3 with differently labeled *CLN3* added via delayed addition I or simultaneous addition (d). (e,f) Confocal images of Whi3 with differently labeled *BNI1* added via delayed addition (e) or simultaneous addition (f). (g,h) Confocal images of Whi3 with differently labeled *BNI1mut* added via delayed addition (g) or simultaneous addition (h). (i) Pearson *r*- values of the colocalization of the differently labeled RNAs. Delayed addition is shown as red bars, whereas simultaneous addition is shown in purple bars. Error bars denote the standard error of mean.

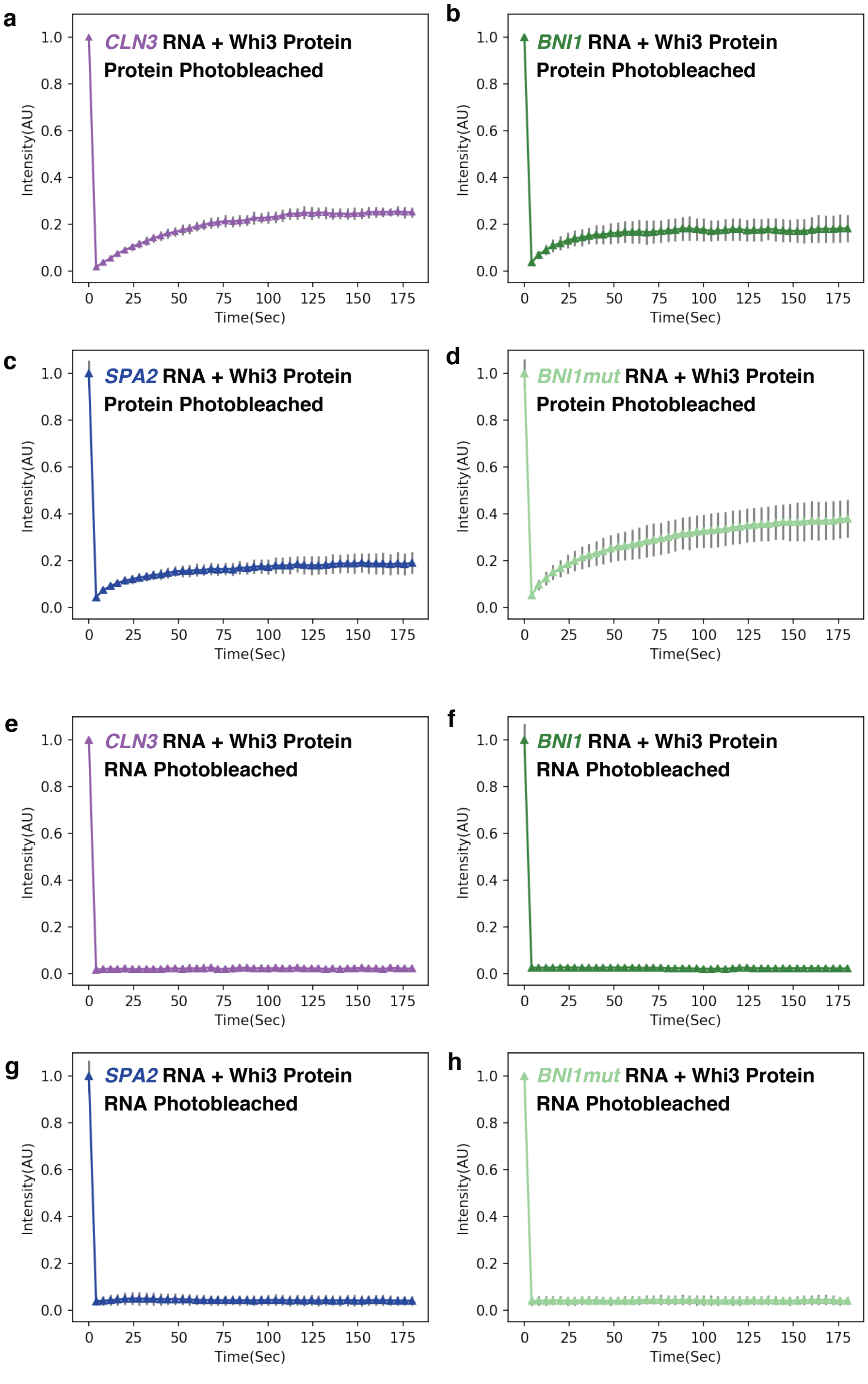


**Figure S11: FRAP traces for two-component Whi3-RNA systems.** (a-d) Fluorescence recovery after photobleaching (FRAP) of Whi3 protein in two-component mixtures consisting of 5 μM Alexa 488-tagged Whi3 protein and 5 nM of the indicated RNA. (e-h) Fluorescence recovery after photobleaching (FRAP) of Cy3-tagged (e) *CLN3* RNA, (f) *BNI1* RNA, (g) *SPA2* RNA, or (h) *BNI1mut* RNA in two-component mixtures consisting of 5 μM untagged Whi3 protein and 5 nM of the indicated tagged RNA. Intensity values are averages of three independent FRAP events with standard error of mean shown as grey bars. Intensity is reported in arbitrary units (AU) normalized to the pre-bleach value.

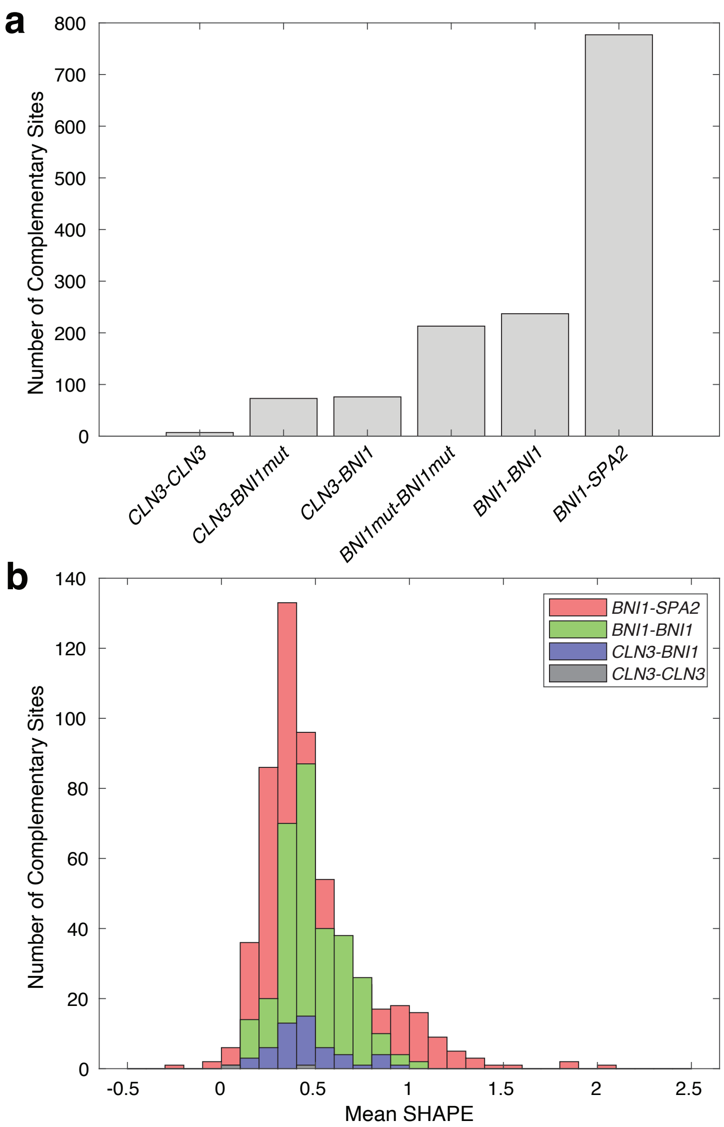


**Figure S12: Number and accessibility of complementary sites between two RNAs.** (a) Number of GUUGle identified complementary sites for different RNA pairs. (b) Histogram of mean SHAPE values for all nucleotides in both complementary sites. Larger SHAPE values imply those complementary sites are more accessible for intermolecular binding.

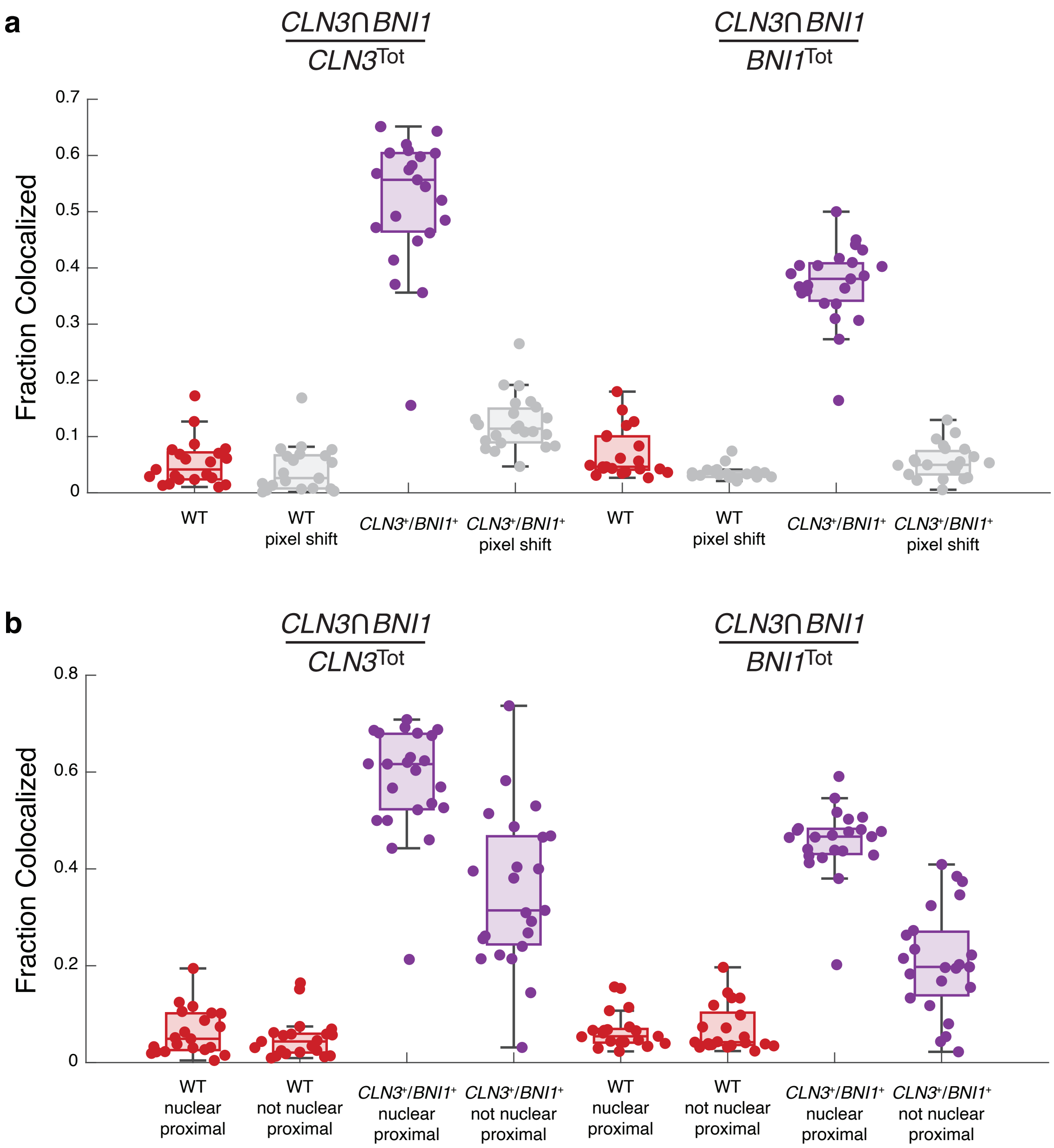


**Figure S13: Pixel shift and subcellular localization analysis of *CLN3* and *BNI1* colocalization.** (a) Fraction colocalized in wildtype (WT) and $CLN3^{+}/BNI1^{+}$ cells with pixel shift data. Here, either *CLN3* or *BNI1* identified spots were shifted by $2r$ in both the x and y direction, where $r$ is the radius of the spot (see Methods). (b) Fraction colocalized based on nuclear proximity. Spots within $r$+$R$ away from the center of any nucleus were defined to be nuclear proximal, where $R$ is the radius of the nucleus (see Methods).
